# Lateral Hypothalamic GABAergic neurons encode and potentiate sucrose’s palatability

**DOI:** 10.1101/2020.06.30.180646

**Authors:** Aketzali Garcia, Alam Coss, Jorge Luis-Islas, Liliana Puron-Sierra, Monica Luna, Miguel Villavicencio, Ranier Gutierrez

## Abstract

Sucrose is attractive to most species in the animal kingdom, not only because it induces a sweet taste sensation but also for its positive palatability (i.e., oromotor responses elicited by increasing sucrose concentrations). Although palatability is such an important sensory attribute, it is currently unknown which cell types encode and modulate sucrose’s palatability. Studies in mice have shown that activation of GABAergic LHA^Vgat+^ neurons evokes voracious eating; however, it is not known whether these neurons would be driving consumption by increasing palatability. Using optrode recordings, we measured sucrose’s palatability while VGAT-ChR2 transgenic mice performed a brief access sucrose test. We found a subpopulation of LHA^Vgat+^ neurons encodes palatability by increasing (or decreasing) their activity as a function of the increment in licking responses evoked by sucrose concentrations. Optogenetic gain of function experiments, where mice were able to choose among available water, 3%, or 18% sucrose solution, uncovered that opto-stimulation of LHA^Vgat+^ neurons consistently promoted higher intake of the most palatable stimulus (18% sucrose). In contrast, if they self-stimulated near the less palatable stimulus, some VGAT-ChR2 mice preferred water over 18% sucrose. Unexpectedly, activation of LHA^Vgat+^ neurons increased quinine intake but only during water deprivation, since in sated animals, they failed to promote quinine intake or tolerate an aversive stimulus. Conversely, these neurons promoted overconsumption of sucrose when it was the nearest stimulus. Also, experiments with solid foods further confirmed that these neurons increased food interaction time with the most palatable food available. We conclude that LHA^Vgat+^ neurons increase the drive to consume, but it is potentiated by the palatability and proximity of the tastant.

## Introduction

The Lateral Hypothalamic Area (LHA) has been regarded as the “feeding center” since its lesion results in hypophagia and subsequent death (Anand and Brobeck, 1951; Teitelbaum and Epstein, 1962). It is part of a neural circuit related to feeding and reward (Delgado and Anand, 1953; Olds and Milner, 1954) as rats are willing to press a lever to deliver electrical intra-cranial self-stimulation (ICSs), and if food is available, it also promotes feeding (Delgado and Anand, 1953; Mendelson, 1967; Mogenson and Stevenson, 1967; Coons and Cruce, 1968). Moreover, if a sweet tastant is available, the rate of electrical ICSs is further increased, whereas bitter compounds decreased them (Phillips and Mogenson, 1968; Poschel, 1968), suggesting an interaction between ICSs and taste palatability. In this regard, and because of its connections with different cortical and subcortical gustatory regions (Simerly, 2004; Berthoud and Münzberg, 2011), the LHA is anatomically located to receive, process, and broadcast taste palatability information (Ferssiwi et al., 1987; Berridge and Valenstein, 1991). Pioneering electrophysiological studies have recorded gustatory-evoked responses in the LHA (Schwartzbaum, 1988; Yamamoto et al., 1989). One recent and elegant study uncovered two functional populations of LHA neurons: one activated by palatable tastants, e.g., sucrose, and another by aversive tastants, like quinine (Li et al., 2013). However, the genetic identity of the LHA cell-type(s) involved in processing palatability-related information remains elusive, and to unveil their identity is a goal of this study.

The LHA is currently viewed as a hub of various cell-types (Stuber and Wise, 2016), grossly divided into two larger populations related to feeding: the glutamatergic (LHA^Vglut2+^) and GABAergic neurons (LHA^Vgat+^) (Gutierrez et al., 2020). Activation of LHA^Vglut2+^ neurons leads to reduced food intake and is aversive (Jennings et al., 2013). In contrast, stimulation of LHA^Vgat+^ neurons is rewarding and produces voracious eating of both foods with nutritional value (Jennings et al., 2015; Navarro et al., 2016) and those without calories, even gnawing behavior towards the cork, an irrelevant biological stimulus (Navarro et al., 2016). Moreover, LHA^Vgat+^ neurons enhance the salience of the nearest stimulus and induce reward via stimulation of their projections to the ventral tegmental area (VTA) (Nieh et al., 2016). In addition, the evoked feeding is mediated by the modulation of terminals reaching a region adjacent to the locus coeruleus (Marino et al., 2020). On the contrary, inhibition of GABAergic LHA^Vgat+^ cell somas is aversive and stops feeding (Jennings et al., 2015; Navarro et al., 2016). However, the role of LHA^Vgat+^ neurons in processing sucrose’s palatability remains to be determined.

This study identified a new role of LHA^Vgat+^ neurons in encoding and enhancing sucrose’s oromotor-palatability responses. Sucrose’s palatability was defined as the enhancement of hedonically positive oromotor responses triggered by increasing sucrose concentrations (Berridge and Grill, 1983; Spector et al., 1998; Villavicencio et al., 2018). Specifically, oromotor responses include an increase in the lick rate and bout size. Thus, it should not be confused with a conscious hedonic feeling of pleasant taste that humans experience (Grill and Berridge KC, 1985; Sclafani, 1991; Berridge and Kringelbach, 2008). We found that opto-stimulation of LHA GABAergic neurons increases the consumption of the most palatable and proximal tastant. They accomplish this by potentiating the palatability of nearby gustatory stimuli. For aversive stimuli, the effect of these neurons is different. In a single bottle test, we found that water deprivation increased the tolerance for bitter compounds and gated a window of opportunity where the activation of GABAergic neurons is sufficient to temporarily reassign the negative hedonic value of quinine and promote its intake. Nevertheless, in a three-option preference test, mice failed to develop a quinine preference when sucrose was available under these neurons’ activation, thus, demonstrating that activation of GABAergic neurons does not merely trigger indiscriminate oromotor tongue movements; rather, the animals’ evoked consummatory behavior largely depends on their internal state and on the palatability of the stimulus. Moreover, optogenetic activation of LHA^Vgat+^ neurons evoked many hallmark behaviors resembling those seen in LHA electrical stimulation. In this regard, and consistent with their positive role in palatability, our results could indirectly explain why electrical ICSs are further facilitated by the presence of sweet solutions (Phillips and Mogenson, 1968; Poschel, 1968). We also found that LHA^Vgat+^ neurons are the common neural substrate for both reward and feeding since after repeated self-stimulation, the more the VGAT-ChR2 mice opto-self-stimulated, the stronger the laser-induced licking they exhibited. We conclude that a subpopulation of GABAergic LHA^Vgat+^ neurons combines stimulus proximity and palatability-related information to enhance nearby energy-rich foods’ palatability and further increase consummatory behaviors.

## Materials and methods

### Animal subjects

We used 42 VGAT-ChR2-EYFP mice (number ID 014548; Jackson Laboratory, Sacramento, CA, USA) and 25 wildtype (WT) littermates, served as controls. Mice were from both sexes between 8-16 weeks old, and they were individually housed in their home cages and maintained in a temperature-controlled (22 ± 1° C) room with 12:12 h light-dark cycle. Unless otherwise mentioned, chow food (PicoLab Rodent Diet 20, MO, USA) and water was available *ad libitum*. For experiments with water restriction, after each behavioral session, mice were allowed to drink water for 1 h daily. All procedures were performed with the approval of the CINVESTAV Animal Care and Use Committee. One session *per* day was conducted between 11:00 a.m. to 2:00 p.m.

### Surgical procedures

All mice were anesthetized with ketamine (100 mg/kg, i.p.) and xylazine (8 mg/kg, i.p.), and then placed into a stereotaxic apparatus (Stoelting, IL, USA). Lidocaine (0.05 ml) was administered subcutaneously under the head’s skin as a local analgesic, ophthalmic ointment (hydrocortisone, neomycin, and polymyxin-B) was applied periodically to maintain eye lubrication. The antibiotic enrofloxacin (0.4 ml/kg) was injected for 3 days after surgery.

For experiments with opto-stimulation of LHA^Vgat+^ cell somas, a single multimode optical fiber 200-μm core diameter with a 0.39 NA (FT200UMT; Thorlabs, NJ, USA) was implanted unilaterally targeting the LHA using the following coordinates: AP: −1.25 mm, ML: ± 1 mm, DV: −4.9 mm, from bregma. For electrophysiology recordings, a custom-made optrode was unilaterally implanted counterbalanced across hemispheres (AP: −1.25 mm, ML: ± 1 mm, from bregma; DV: −5.2 mm ventral to dura). The optrode comprises an array of 16 tungsten wires formvar coated (35 μm diameter) surrounding and protruding 1 mm the single multimode optical fiber tip. No significant differences were found between hemispheres, so data were pooled together (data not shown). All experiments began 7 days after surgery to allow recovery.

### Histology and immunofluorescence

Mice were sacrificed by an overdose of pentobarbital (150 mg/kg) and perfused with PBS, followed by 4% paraformaldehyde. Brains were fixed overnight in 4% paraformaldehyde and gradually replaced in a gradient of concentrations until 30% sucrose. For histology, brain slides (40 μm) were cut with a cryostat (Thermo Scientific HM525 NX), and images were taken with a Nikon eclipse e200 microscope and a Progress Gryphax microscope camera, using a 10x objective. For immunofluorescence, free-floating brain slides (40 μm) were blocked with 1% BSA, 0.2% Triton in PBS for 30 min. They were then washed with 0.2% Triton in PBS three times every 10 min, followed by the incubation with the primary antibodies: mouse anti-GAD 67 primary antibody (Millipore, Mab5406, 1:1000 dilution), and rabbit anti-GFP primary antibody (Invitrogen, A11122, 1:1000 dilution). Incubation took place overnight at 4°C. The next day, brain slides were washed with 0.2% Triton in PBS three times every 10 min, then incubated with the secondary antibodies Alexa 647 goat anti-mouse (Invitrogen, A21235, 1:500 dilution), and Alexa 488 donkey anti-rabbit (Invitrogen, A21206, 1:500 dilution) during 90 min at room temperature. Afterward, we applied DAPI to stain the nuclei. Brain slides were put on slides with a mounting medium for fluorescence (Vectashield), and images were taken with a Leica confocal microscope using a 63x objective.

### Gustatory stimuli

Sucrose reagent-grade chemical quality (Sigma-Aldrich, Mexico City, Mexico) was used in the following concentrations: water, 3, 10, and 18 wt%. Quinine hydrochloride dihydrate (QHCl) reagent-grade at 0.04 wt% (Sigma-Aldrich, Mexico City, Mexico). Artificial Saliva contained (mM): 4 NaCl, 10 KCl, 6 NaHCO3, 6 KHCO3, 0.5 CaCl2, 0.5 MgCl2, 0.24 K2HPO4, 0.24 KH2PO4 (Zocchi et al., 2017). We added 0.05 mM HCl to adjust to pH 7. The solutions were dissolved in distilled water, maintained under refrigeration, and used at room temperature. We also used 20 mg chocolate pellets (Bio-Serv, NJ, USA), Chow food (PicoLab Rodent Diet 20, MO, USA), a high-fat diet with 45% kcal% fat (Research Diets, NJ, USA), granulated sugar cube, and cork.

### Electrophysiology

Neural activity was recorded using a Multichannel Acquisition Processor System (Plexon, Dallas TX, USA) interfaced with Med Associates to simultaneously record behavioral events (Gutierrez et al., 2010). Extracellular voltage signals were first amplified by an analog head-stage (Plexon, HST/16o25-GEN2-18P-2GP-G1), then amplified and sampled at 40 kHz. Raw signals were band-pass filtered from 154 Hz to 8.8 kHz and digitalized at 12 bits resolution. Only single neurons with action potentials with a signal-to-noise ratio of 3:1 were analyzed (Gutierrez et al., 2010). Action potentials were isolated online using a voltage-time threshold window, and three principals’ components contour templates algorithm. Furthermore, off-line spike sorting was performed (Plexon, offline sorter), and only single units with stable waveforms across the session were included in the analyses (Gutierrez et al., 2010); see Supplementary Figure 1. Also, to verify waveform stability, we correlated waveform’s shapes recorded in the brief access test and the optotagging session.

### Optogenetic stimulation

A 473 nm laser intensity was modulated by a DPSS system (OEM laser, UT, USA). Laser pulses were synchronized with behavioral events with Med Associates Inc. software and TTL signal generator (Med Associates Inc., VT, USA). The patch cord’s optical power was at 15 mW, and it was measured with an optical power meter (PM20A, Thorlabs, NJ, USA). However, we delivered between 10 to 12.6 mW at the fiber optic tip, depending on each fiber’s efficiency. Unless otherwise mentioned, the laser was turned on by 2 s (at 50 Hz) and 4 s off, with 10 ms pulse width and a duty cycle of 50%.

### Stimulation’s parameters for LHA GABAergic neurons

To explore the best stimulation parameters for LHA GABAergic neurons, mice were implanted with an optrode in LHA and placed in a custom-made box with 18 x 13 x 7.5 cm. With no food available, the laser was turned on at different frequencies 0, 2.5, 5, 10, 20, and 50 Hz semi-randomly, while animals’ neural activity was recorded for 20 min. We used a Kruskal-Wallis test to compare firing rates during the baseline (from −1 to −0.04 s) against the activity during laser presentation (from 0 to 2 s) aligned (time = 0 s) to laser onset for all frequency tested. Neurons that significantly increased their firing rate during the laser stimulation were named “Activated,” and neurons that decreased their activity were named “Inhibited.”

### Neural activity recording during a brief access test and palatability-related responses

We used a brief access test in water-deprived mice while the LHA activity was recorded for 30 min. We employed a licking spout consisting of independent stainless-steel needles (20-gauge) cemented together with dental acrylic at the sipper tube’s tip (Villavicencio et al., 2018). Each needle was connected to a solenoid valve (Parker, Ohio, USA) by a silicon tube. The drop (~2 μL) was calibrated before starting the session, using a pressurized control system. The trial structure was as follows: At the beginning of the task, the sipper was extended in lick position. To start a new trial, each mouse was required to introduce its head into the central port and then elicited a dry lick to start the reward epoch (7 s). During this period, a tastant’s drop was delivered every lick. The sipper was retracted for 3 s as Inter-Trial Interval (ITI), and then re-extended in a lick position into the central port. Tastant solutions (artificial saliva, water, sucrose 3%, and sucrose 18%) were delivered in a semi-random order.

To identify neurons whose firing rate correlated with palatability-induced oromotor responses, we used a best-window analysis based on a previous study (Villavicencio et al., 2018). This method has been used to detect palatability-related activity in a brief access test. Briefly, during each recording session, a palatability-index (PI) was calculated. The PI reflects the overall appetitive oromotor responses elicited by each tastant delivered in the session. It was computed by averaging the lick rate during the entire reward epoch, including all the trials per session. In mice, the PI takes values between 0-8 Hz for sucrose stimuli (Glendinning et al., 2005). 0 Hz means animals ultimately rejected a solution in all trials, whereas 8 Hz indicates they licked continuously during the entire reward epoch, thus reflecting a greater palatability response elicited by the tastant. Then, the firing rate was calculated for a variety of time centers (from 0.25 to 6.75 s with 0.5 s steps) and multiple window sizes (from 0.5 to 7 s, 0.5s steps), such that each window was estimated as the center ± (window size/2). We identified the windows where the mean firing rate was significantly different for at least one tastant delivered (i.e., tastants, using a Kruskal-Wallis test at an alpha of 0.05). Next, we computed Pearson’s correlation coefficient (r; the alpha level at 0.05) between both the PI and the firing rate on a trial-by-trial basis. The window with the largest absolute Pearson’s correlation coefficient was selected as the “best-window.” Thus, for the “best-window” and each statistical test (i.e., Kruskal-Wallis test and Pearson’s correlation coefficient), a permutation assay was used for multiple-testing correction. This analysis was accomplished by shuffling the different tastant delivered 20,000 times (using the Matlab function “shuffle”). A corrected *p*-value was obtained using the following formula *p*= (k+1)/(n+1), where k is when a permuted distribution leads to a *p*-value smaller than the original *p*-value and n is the number of repetitions. Only time-windows with *p*’s < 0.05 in both tests (Kruskal-Wallis and Pearson Correlation) were considered statistically significant. Thus, the “best-window” is where the firing rate maximally correlated with the oromotor responses (the lick rate) elicited by sucrose’s palatability on a trial-by-trial basis. Importantly, results were qualitatively similar if we used the lick bout size rather than the lick rate to compute the PI. A lick bout was defined as the collection of at least two rhythmic licks separated by a pause ≥ 500 ms during the reward epoch. The bout size was the time difference (in seconds) between the last and first lick in the bout (Gutierrez et al., 2006).

To evaluate whether palatability-related neurons dynamically track the changes in lick rate across the session, we computed the lick rate in the reward epoch as a function of trial types (artificial saliva (AS), water, sucrose 3%, and sucrose 18%), divided into blocks of 10^th^ percentile of trials each. We verified that every block across the session has the same number of trials.

### Optotagging task

Once the brief access test was finished, each mouse was tested in the optotagging task. For this task, we removed the licking spout from the behavioral box. Over the session (15 min), the laser was turned on (at 50 Hz) during 2s or 7 s, followed by 10 s off. Laser activated neurons (pLHA^Vgat^) were detected by using two methods: 1) a Kruskal-Wallis to compare firing rates during a baseline (from −1 to −0.04 s) against the activity during the presentation of laser (from 0 – 2 s, aligned to laser onset). 2) Cumsum statistic (Gutierrez et al., 2006) to obtain the first bin (1 ms resolution) after the first laser pulse that significantly increased the firing rate above baseline (from −20 to 0 ms). Only neurons showing a significant value in both tests were considered laser-activated pLHA^Vgat^ From the population pLHA^Vgat^ neurons, we identified two types of populations: 1) ChR2-expressing cells LHA^Vgat+^ or early neurons. These neurons were those with an action potential evoked within ≤ 15 ms latency (Buonomano, 2003), measured by a cumsum statistic; and 2) laser-activated late neurons: these were those with an action potential occurring after > 15 ms latency. Also, to classify non-LHA^Vgat^ neurons, we used Kruskal-Wallis test as described above. Neurons with no significant modulation were classified as “unmodulated.”

### Open loop stimulation with one option

The front panel of an operant conditioning chamber (Med Associates Inc, VT, USA) was equipped with a central port and a sipper, where individual licks were registered by a contact lickometer (Med Associates Inc, VT, USA). To determine the best stimulation parameters to induce feeding behavior, a group of naïve mice had free access to sucrose 10% solution. For open loop stimulation, mice were opto-stimulated by alternating blocks of 5 min “off” and 5 min “on” across a 40 min session (Nieh et al., 2016). During opto-stimulated blocks, the laser was turned “on” regardless of the mice’s behavior and spatial position in the chamber. A different opto-stimulation frequency (0, 2.5, 5, 10, 20, and 50 Hz) was delivered daily. The laser bound feeding index was measured as the number of licks in a 2.5 s window from laser onset divided by total licks in the session, hereafter named laser bound feeding, and it was plotted as a function of laser frequency.

### Pre-stimulation task

A group of naïve mice was placed in an operant conditioning chamber with a central port. A door blocked the access to the sipper during the first 15 min (pre-stimulation period. Each mouse was opto-stimulated with all pre-stimulation protocols (0, 5, 10, and 15 min) following a Latin square design. Then, the door was opened, and a 10% sucrose solution was available during the next 15 min. During the pre-stimulation period, the laser was turned on for 2 s and off for 4 s.

### Open loop stimulation with three options

To determine whether opto-stimulation of LHA GABAergic neurons drives the intake with a bias towards the most palatable stimulus available, a new group of naïve mice was tested in an open loop stimulation task (alternating 5 min no-laser and 5 min “on” (2 s on, 4 s off) blocks across a 40 min session). The operant chamber was equipped with three ports, where mice had free access to water (central sipper), a 3% sucrose solution (left sipper), and 18% sucrose solution (right sipper). Lateral sippers were counterbalanced across animals. This task comprises four baseline sessions (with no photostimulation, data not shown), and eleven test sessions were pooled together. The number of licks given to the sipper filled with sucrose 18%, across the eleven sessions was also plotted.

For this experiment, we video tracked the mouse’s distance and head angle relative to each of the three sippers. The sippers’ position and three anatomical animal points, the nose, the neck, and the tail’s base, were extracted using the DeepLabCut Python package (Mathis et al., 2018; Nath et al., 2019). We define the mouse position as the coordinates (x and y) for the nose, whereas the head direction was a 2D vector going from the neck to the nose. For each video frame, each sipper’s distance was calculated (in pixels), and the angle between the head direction and position of each sipper (in degrees). An angle of 0° means the mouse is looking at the sipper, and 180° means facing the opposite direction. For distance measuring, 100 pixels correspond to approximately 6 cm. The mouse position and direction were extracted from the train of stimulation’s first laser pulse, and the following 6 s window was analyzed to find out if the mice initiated a licking behavior at any of the 3 sippers. Then, we built a two-dimensional array containing the probability of licking any sipper given the mouse’s distance and angle during the first laser of each opto-stimulation train.

### Open loop stimulation in an open field with Chow, high-fat diet, or granulated sugar cube

A new group of animals was placed in the circular arena’s center (50 cm in diameter). Three or two food plates were equidistant to each other, and each contained either Chow, high-fat diet, or granulated sugar cube. Mice were opto-stimulated by alternating 5 min block with no-laser and 5 min laser block across the session (20 min). In open loop, the laser was turned “on” regardless of the mice’s behavior and spatial position in the arena. All sessions were recorded and analyzed with bonsai software (https://open-ephys.org/bonsai). We calculated the mouse’s centroid at each frame and used that information to create a heatmap of its position. The distance of the mouse’s centroid from each stimulus was computed. A food interaction occurred when the distance was below 60 mm, and the animal stayed there for at least 1 s. Food plates were weighed at the beginning and the end of each session.

### Closed-loop stimulation with three options

To evaluate whether activation of LHA^Vgat+^ neurons induces preference for the most proximal stimulus, we used a closed-loop stimulation protocol (in the same mice from Open loop stimulation with three options), over eleven additional sessions (40 min each). As noted above, mice were placed in an operant conditioning chamber with three ports (water, central sipper; 3% sucrose solution, left sipper; 18% sucrose solution, right sipper). Photostimulation was delivered when subject made a head entry in the central port (containing a sipper filled with water), laser was turned on by 2 s followed by a time out of 4 s (where the laser could not be re-activated), after this, a new laser onset (2 s “on”) occurred when mice performed a new head-entry. Thus, in this experiment, the less palatable stimuli (water) was the nearest to opto-self-stimulation. During photostimulation, sucrose preference index was measured as the number of licks of 18% sucrose divided by total licks for sucrose 18% + water. Values higher than 0.5 indicate sucrose preference.

To evaluate the laser-bound feeding development across the sessions, we correlated laser bound feeding during open loop stimulation sessions vs. closed-loop stimulation for the first three and the last five sessions.

### Closed-loop in an open-field

In a circular arena, four stimuli were located equidistant to each other: wood cork, Chow, chocolate pellets, or a sipper filled with a 10% sucrose solution. During the session, a homemade computer vision program tracked in real-time the position of the subject. Each mouse needs to enter the designated area to receive opto-self-stimulation (2 s laser on and 4 s off). Mice need to leave and re-enter the designated area to begin a new trial. Only one designated area was used per session, and it remained in the same position for up to 3 or 4 consecutive sessions. Each session’s duration was 40 min (mice were the same as those used in the previous task).

### Closed-loop stimulation with different options in the central port

For the same group of mice used in the Closed-loop with three options, and on subsequent days, the central port’s stimulus was replaced across twenty-eight sessions, following the protocols described in Table 1.

**Table 1.** Protocol used for closed-loop stimulation with central port stimulus replaced.

| Stimuli in central port | Sessions tested |
| --- | --- |
| Water | Last 5 sessions of the closed-loop |
| Extinction (No opto-stimulation) | 4 |
| Water | 3 |
| Empty sipper | 3 |
| Water | 3 |
| 0.04% Quinine | 4 |
| Airpuff | 4 |
| Unavoidable airpuff | 3 |
| Sucrose 18% (Water was placed in the sucrose 18% port) | 4 |

### Closed-loop stimulation task with one option

Mice were put in a smaller and custom-made box with 18 x 13 x 7.5 cm internal dimensions. The front wall was equipped with one single sipper and a contact lickometer with a V-shape to register individual licks. First, mice were tested, for 3 days, with a 0.04% quinine solution under sated state (fed mice) and then for 3 more days under a 23 h water-deprived condition, each session lasted 20-min. Then, mice were tested for 2 days with a sipper filled with 18% sucrose, in a sated state, and then 2 more days with water-deprived condition. Opto-self-stimulation (2 s on with a time out of 4 s, where the laser could not be activated) was delivered when mice made a head entry in the central port. Water-deprived mice had access to *ad libitum* water for 1 h after the session.

### Opto-self-stimulation during a brief access test

To evaluate whether activation of GABAergic neurons enhances oromotor-palatability responses, we performed a brief access test with a new group of mice over 31 sessions. The behavioral setup conditions were the same as described above, and as tastant solutions, we used water, sucrose 3%, and sucrose 18% (delivered in a semi-random order). The trial structure was similar, but this time to start a new trial, each mouse was required to introduce its head into the central port to turn “on” the laser. Then, elicited a dry lick to start the reward epoch (7 s). Each head entry triggers the laser onset for 2s “on” (or in different sessions for 7 s “on”; see Table 2). At the end of the session, mice had access to *ad libitum* water for 1 h. Protocols are described in Table 2 below:

**Table 2.** Protocol for opto-self-stimulation in a brief access test.

| Laser on | Time on | # Sessions<br>(30 min each) |
| --- | --- | --- |
| All trials | 2 s | 6 |
| Water trials | 2 s | 3 |
| Sucrose 3% trials | 2 s | 3 |
| Sucrose 18% trials | 2 s | 3 |
| All trials | 7 s | 4 |
| Water trials | 7 s | 3 |
| Sucrose 3% trials | 7 s | 3 |
| Sucrose 18% trials | 7 s | 3 |
| Mock laser (sucrose 3% trials) * | 7 s | 3 |

\*For mock laser sessions, mice were connected to a false fiber optic, whereas the real laser was glued outside the skull to emit blue light.

In addition, to verify that LHA GABAergic neurons were capable of sustained activation over 7 s of optostimulation. We plotted the PSTH of pLHA^Vgat^, non-LHA^Vgat^, and unmodulated neurons using the same method previously described.

### Data Analysis

All data analysis was performed using MATLAB (The MathWorks Inc., Natick, MA) and GraphPad Prism (La Jolla, CA, USA). We used the mean ± sem and the α level at 0.05.

## Results

Initially, we determined the optimal stimulation parameters to activate LHA GABAergic neurons. This was accomplished by implanting an optrode in naïve mice that constitutively expressed ChR2, fused with an enhanced yellow fluorescent protein (EYFP) in GABAergic neurons expressing the gene for the vesicular γ-aminobutyric acid transporter (hereafter referred to as VGAT-ChR2 mice) (Zhao et al., 2011) (Figures 1A-B). We found that a laser stimulation of 50 Hz (10 ms width) evoked the strongest LHA neuronal responses (Figures 1C-F) (Gigante et al., 2016). Unless otherwise mentioned, this frequency was used for all subsequent experiments.

**Figure 1.**
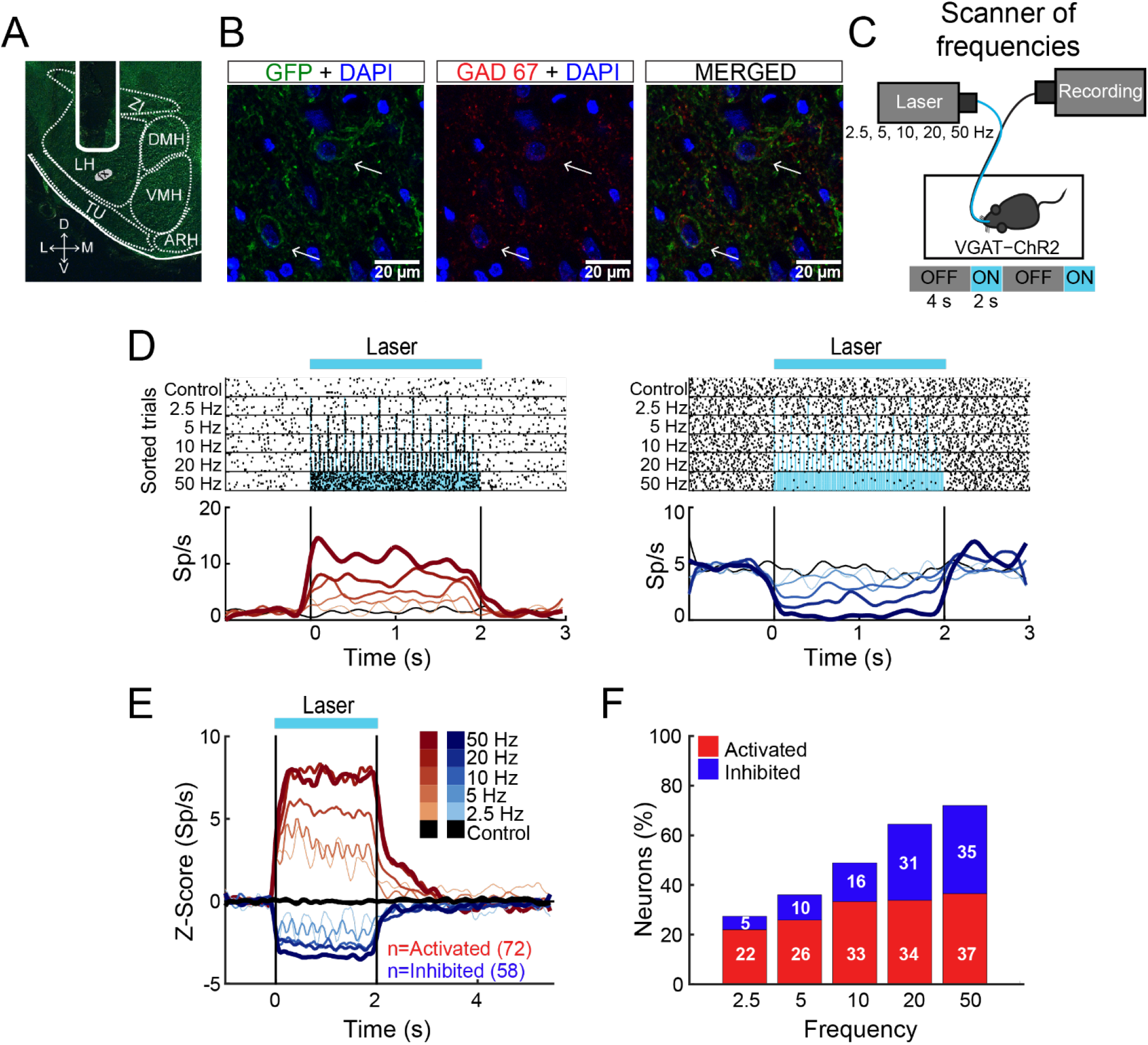
The 50 Hz stimulation evoked the largest LHA neuronal responses. **(A)** Representative location of an optrode in LHA of VGAT-ChR2 mice. **(B)** Confocal images from LHA immunostained for GFP (green), GAD67 (red), and DAPI (blue). Arrows indicate the colocalization of ChR2-EYFP and GAD67 (see merged). **(C)** Schematic of optrode recordings in LHA during control trials (with no-laser) and opto-stimulation at 2.5, 5, 10, 20, and 50 Hz delivered randomly. In each trial, the laser had a cycle of 2 s on and 4 s off. **(D)** Representative raster plots of two neurons recorded in LHA. The first one was activated (left panel), and the second inhibited (right panel) during opto-stimulation. Spiking responses were aligned (Time = 0 s) to the first laser pulse. Black ticks depict action potentials and blue marks laser pulses. Below are the corresponding PSTHs (firing rates, Sp/s). Red and blue ascending gradients indicate the frequency of stimulation (see E for the color bar). Black trace represents activity during control trials without laser. Vertical black lines indicate laser onset (Time = 0 s) and offset (Time = 2 s), respectively. **(E)** Normalized Z-score population responses of LHA neurons recorded from n = 5 VGAT-ChR2 mice. Red and blue colors depict activated or inhibited responses, respectively, relative to baseline (−1 to 0 s), thickness, and gradient colors indicate higher stimulation frequencies. Black trace illustrates neural activity during control trials for all recorded neurons (n = 186). **(F)** Percentage of neurons recruited as a function of stimulation frequency.

### LHA neurons encode sucrose’s oromotor palatability responses

In rats, previous electrophysiological studies suggest that ensembles of LHA neurons process palatability-related information (Norgren, 1970; Schwartzbaum, 1988; Yamamoto et al., 1989; Li et al., 2013). Based on these results, we then asked whether LHA neurons could provide information about sucrose’s palatability using a behavioral task with only palatable stimuli (i.e., artificial saliva (AS), water, sucrose 3 wt%, and sucrose 18 wt%). To this end, taste responses were recorded from a total of 284 LHA neurons, while VGAT-ChR2 mice performed a brief access test (Figure 2A). To uncover palatability-responsive neurons, we analyzed neuronal activity using a previously described “best-window analysis” (Villavicencio et al., 2018) (see Materials and Methods). This strategy detects the optimal interval, where the firing rate best correlates with the sucrose’s palatability index (PI) computed as the average lick rate (or bout size), on a session by session basis, evoked by each gustatory stimulus in the Reward epoch (Figure 2B; one-way ANOVA, F_(3, 108)_ = 10.35, *p* < 0.0001; Supplementary Figure 2A; one-way ANOVA, F_(3, 108)_ = 2.926, *p* < 0.05). Figure 2C depicts the responses of two representative LHA neurons: as expected, we found one whose activity increased as the sucrose concentration increased (positive Pearson’s correlation, r = 0.87; Figure 2C *left*), and the other was negatively associated with palatability oromotor responses evoked by sucrose (r = −0.64; Figure 2C *right)*. Figure 2D shows the normalized activity exhibiting either positive (59/284 neurons, 21%) or negative correlations (76/284, 27%) with the PI. Also, qualitatively similar results were found using the bout size to compute the PI (Supplementary Figure 2B). Thus, these data confirmed that the LHA mice’s neurons exhibit palatability-related responses, and this activity can be either correlated or anti-correlated with the sucrose solution’s PI.

**Figure 2.**
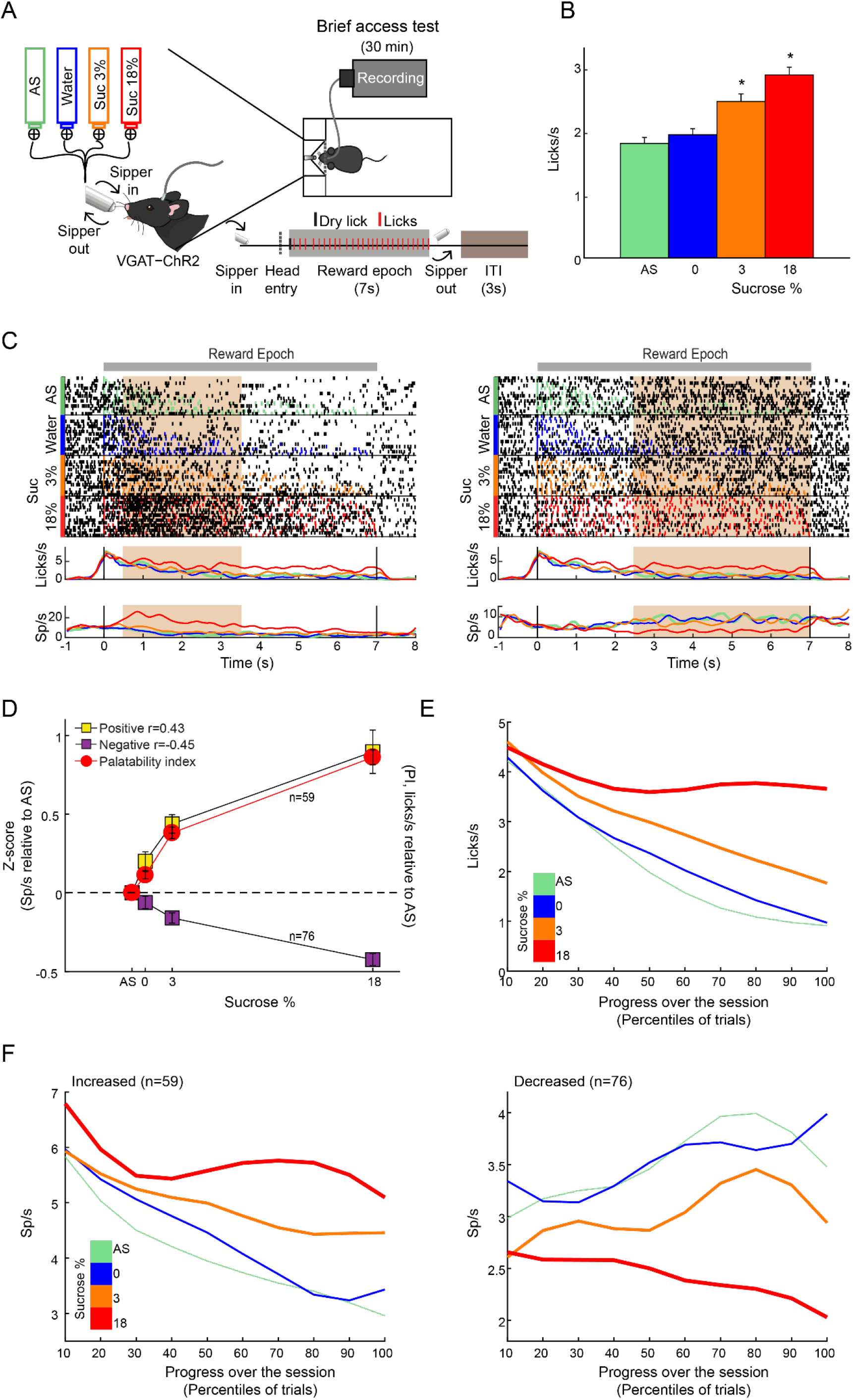
LHA neurons process palatability-related information. **(A)** Brief access taste setup depicting a behavioral box equipped with a computer-controlled retractile sipper that for each lick independently delivers a drop of artificial saliva (AS), water, sucrose 3%, or 18% concentrations. After a head entry (dashed line), the first dry lick (black line) enables the 7 s reward epoch. During this period, in each lick (red), a drop of the four tastants was randomly delivered. After the reward epoch, the sipper was retracted for an Intertrial Interval (ITI) of 3 s and then re-extended to start a new trial. **(B)** Average lick rate during the entire reward epoch, reflecting greater palatability as a function of sucrose concentration. * Indicates significant difference (p < 0.05) among stimuli. Using one-way ANOVA followed by the Holm Sidak test. **(C)** Representative raster plot of two LHA neurons recorded while mice performed a brief access test. Spikes are plotted as black ticks, whereas licks for each tastant are color-coded (green, AS; blue, water; orange, sucrose 3%; red, sucrose 18%). Below are corresponding peristimulus time histograms (PSTHs) of lick rate (Licks/s) and firing rate (Spikes/s; Sp/s) as a function of trial type. Neuronal responses were aligned (Time = 0 s) to the first rewarded lick. The brown rectangles depict the “best” window with the maximum Pearson correlation coefficient between firing rates and sucrose’s palatability index (PI). The left and right raster plots displayed two neurons with a positive and negative correlation, respectively. **(D)** Z-score normalized activity (relative to AS trials) for LHA neurons with either positive or negative correlation against PI (red circles). **(E)** Population PSTHs of the lick rates given in the Reward epoch as a function of trial type, divided into blocks of 10^th^ percentiles of trials each. **(F)** Population PSTHs of the firing rate during the Reward epoch of the palatability-related neurons across the session for each trial type. *Left panel* depicts neurons that fired more to higher sucrose concentrations (positive sucrose’s palatability correlation; (D) yellow), whereas, *right panel* illustrates neurons with decreasing firing rates as the sucrose concentration increased (i.e., negative correlation with sucrose’s palatability; (D) purple).

### Dynamic tracking of palatability over the entire course of a behavioral session

Having demonstrated that LHA neurons encode sucrose’s palatability-induced oromotor responses, we then explored whether LHA palatability-related neurons can dynamically adjust their responses to track the changes in sucrose’s palatability across the session when the animals would approach satiation. We found that the lick rate declined across the session as a function of sucrose’s palatability (Figure 2E; RM ANOVA, main effect of stimuli, F_(3, 402)_ = 163.7, *p* < 0.0001;effect of time, F_(9, 1206)_ = 100.4, *p* < 0.0001; and stimuli by time interaction, F_(27, 3618)_ = 16.54, *p* < 0.0001). Briefly, early in the session, animals licked more for all stimuli, but as the session progressed, the lick rate declined, especially for AS and water but not for sucrose (3% and 18%), indicating that they tracked palatability responses rather than satiety (Figure 2E). Similar to what we have shown in the Nucleus Accumbens Shell (NAcSh) (Villavicencio et al., 2018), a brain region that sends outputs to LHA and is involved in feeding and also contains palatability-related neurons (O’Connor et al., 2015; Villavicencio et al., 2018), we also found that the LHA neuron’s firing rate (either increasing or decreasing neurons) adjust across the session to reflect sucrose’s palatability (Figure 2F). These data indicate that within the session, as the animals approach satiety, the responses of the Palatability related-neurons tracked the rapid decline in the lick rate mainly for the two less palatable stimuli (i.e., AS and water) but not for the higher sucrose-concentration (sucrose 3% and 18%), suggesting these neurons are tracking palatability and no satiety.

### A subpopulation of opto-identified LHAVgat+ neurons encodes sucrose’s palatability

To identify LHA^Vgat+^ neurons from our recordings, we opto-stimulated the same mice recorded in the brief access test seen in Figure 2. We verified the stability of the waveforms between tasks (i.e., brief access test and optotagging). We only included single units with stable waveforms in the analysis (see Materials and Methods; Supplementary Figure 1). Figure 3A displays the setup and the normalized (Z-score) activity of laser-activated neurons (48%; 137/284) (for details, see Material and Methods). Since these neurons may include ChR2 expressing cells and other LHA neurons modulated by indirect polysynaptic feedback of afferent fibers from other areas in the brain (Nieh et al., 2015), we decided to name them putative LHA^Vgat^ (pLHA^Vgat^) neurons. Note that pLHA^Vgat^ neurons comprise the ensemble recruited by the optogenetic stimulation of LHA GABAergic neurons, suggesting they convey similar information. Therefore, we next evaluated whether pLHA^Vgat^ neurons were correlated with sucrose’s palatability. In total, 50% (69/137) of laser-activated neurons exhibited responses that were significantly palatability related; specifically, 34% showed a positive correlation (46 out of 137), whereas 17% (23 out of 137) had a negative correlation with sucrose’s palatability (Figure 3B; Positive vs. Negative chi-square test (1, 274) = 10.25, *p* < 0.01). Figure 3C shows the population activity of pLHA^Vgat^ neurons with either a positive (yellow) or negative (purple) Pearson’s correlation coefficients with the PI (red). Furthermore, from all recorded LHA neurons that encode sucrose induced oromotor-palatability responses with a Positive correlation (n = 59, see Figure 2D), we found that more than 78% (46/59) belonged to the pLHA^Vgat^ (Figure 3D). In contrast, only 30% (23/76) of pLHA^Vgat^ neurons were Negative palatability related (Figure 3D; Positive vs. Negative chi-square test (1, 135) = 30.25, *p* < 0.0001). Similar results were found when we analyzed the Laser-activated late neurons (hereafter named Late neurons) from the pLHA^Vgat^ population. These neurons exhibited a delayed laser evoked action potential with a slow latency (> 15 ms), n=66, see Supplementary Figure 3A. We compared the proportion of Late neurons against all palatability-responsive neurons recorded in the LHA and found that 41% (24/59) belonged to the Positive palatability-related population, and only 8% (6/76) were Negative palatability-related neurons (Supplementary Figure 3B-C; chi-square test (1, 135) = 20.65, *p* < 0.0001). Thus, pLHA^Vgat^ encode sucrose’s palatability with a biased towards positive correlations.

**Figure 3.**
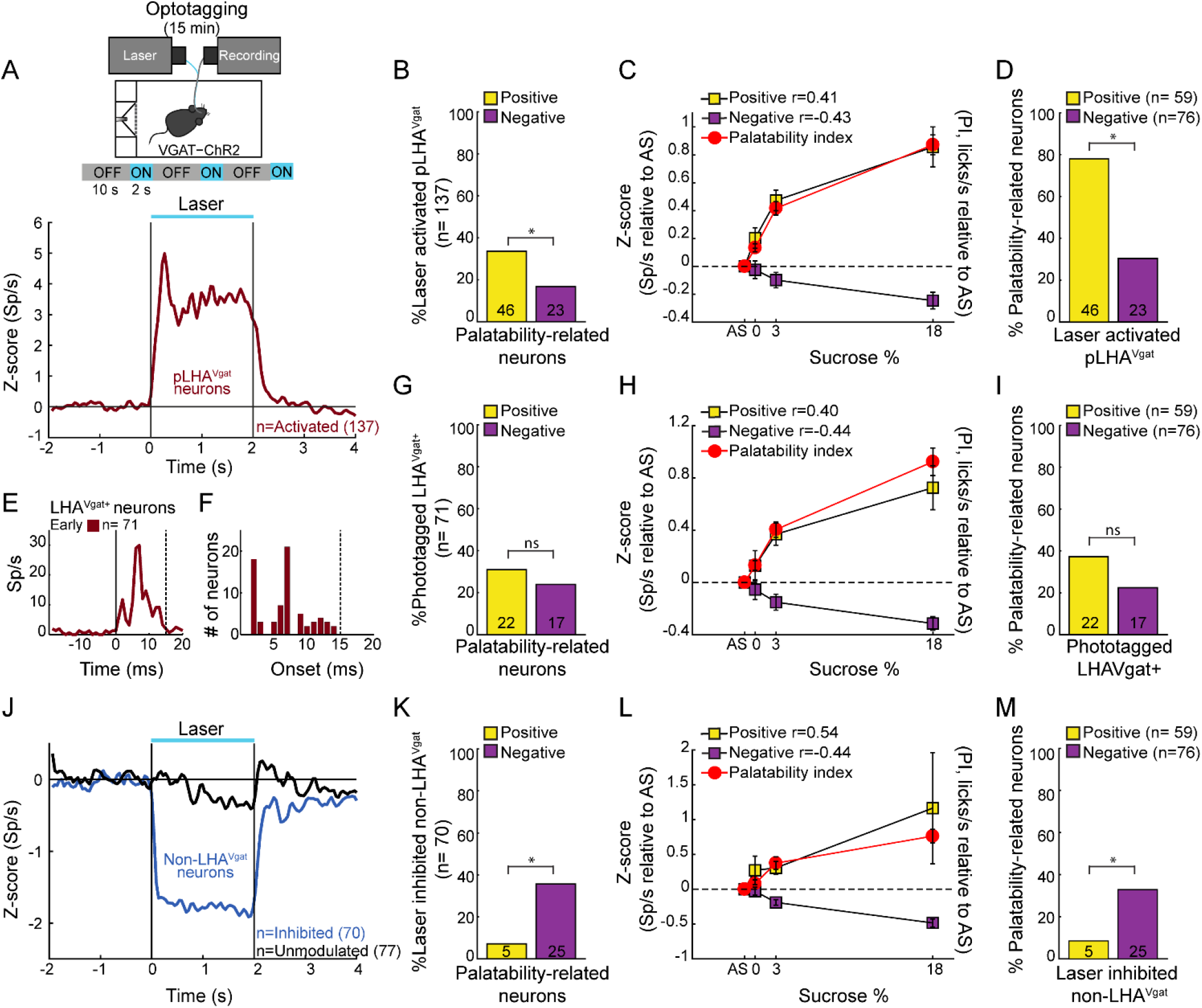
A subpopulation of LHA^Vgat+^ neurons encodes sucrose’s palatability. **(A)** To confirm the LHA^Vgat+^ neurons’ identity after the brief access test, we performed laser opto-stimulation in a cycle 2 s on (at 50 Hz) and 10 s off over 15 min session. In this phase, mice had no access to the sipper (*above panel*). *Below,* the normalized Z-score population activity of all 284 LHA neurons recorded from 3 VGAT-ChR2 mice. Responses were aligned to the laser onset (Time = 0 s), relative to baseline (−1 to 0 s) activity. The red line depicts the responses of activated putative pLHA^Vgat^ neurons. **(B)** Percentage of laser-activated pLHA^Vgat+^ neurons whose activity (in the brief access test) also positively or negatively correlates with sucrose’s PI. * *p* < 0.01, chi-square. **(C)** Z-score activity (relative to AS) for all activated pLHA^Vgat^ neurons and its correlation with the sucrose’s PI. **(D)** Percentage of Positive or Negative laser-activated palatability-related neurons. * *p* < 0.0001, chi-square. **(E)** PSTH of identified (opto-tagged) LHA^Vgat+^ neurons exhibiting early evoked responses (i.e., ChR2 expressing cells). **(F)** Histogram of opto-tagged LHA^Vgat+^ neurons with evoked early responses (latencies < 15 ms) from laser onset. **(G-I)** same conventions as in B-D, but for opto-tagged LHA^Vgat+^ neurons encoding sucrose’s palatability. **(J-M)** Non-LHA^Vgat^ neurons that negatively encode sucrose’s palatability. ***J,*** Blue and black traces correspond to Inhibited (Non-LHA^Vgat+^ neurons) and Unmodulated responses, respectively. Same conventions as in (*A)*. **(K-M)** same conventions as in B-D, * *p* < 0.001, chi-square.

To confirm the identity of LHA^Vgat+^ neurons, we searched for neurons in which a brief pulse of light-evoked an action potential with an early latency (≤15 ms) that would reflect the expression of ChR2 in their somas (Buonomano, 2003). In total, from the pLHA^Vgat^ population, 52 % (71/137) neurons exhibited an early response to blue light (Figures 3E-F), suggesting that these pLHA^Vgat^ neurons were LHA^Vgat+^ neurons. We found that over half of the identified LHA^Vgat+^ neurons, 55% (39/71), were involved in encoding sucrose’s oromotor-palatability, and the remaining (45%) were unmodulated by sucrose. From the LHA neurons related to sucrose’s oromotor-palatability, 31% of LHA^Vgat+^ neurons positively correlated with sucrose’s palatability (22/71), and 24% were anticorrelated (17/71) (Figure 3G; Positive vs. Negative chi-square test (1, 142) = 0.88, p = 0.34). The LHA^Vgat+^ neuronal responses with positive and negative Pearson’s correlation coefficients with lick-related palatability responses are seen in Figure 3H. Once again, we compared the proportion of opto-identified LHA^Vgat+^ neurons against all palatability-responsive neurons recorded in the LHA. We found a trend to encode sucrose’s palatability in a positive rather than negative manner. That is, 37% (22/59) of LHA^Vgat+^ neurons belonged to the Positive palatability-related population, whereas only 22% (17/76) were Negative palatability-related neurons (Figure 3I; Positive vs. Negative chi-square test (1, 135) = 3.59, p = 0.057). Altogether, these data suggest that LHA^Vgat+^ neurons tend to encode sucrose’s palatability in a positive rather than in a negative manner. More important, our data agree with the idea that the LHA^Vgat+^ neurons comprised heterogeneous subpopulations with different functional responses (Jennings et al., 2015).

### Non-LHA^Vgat^ neurons negatively encode sucrose’s palatability

In contrast, we found that 25% of LHA neurons exhibited a laser-induced inhibition (70/284, non-LHA^Vgat^ neurons, Figure 3J, blue trace), and the remaining 27% (77/284) were unmodulated during blue light stimulation (Figure 3J, black trace). Unlike pLHA^Vgat^ neurons, only 7% (5 out of 70) of the non-LHA^Vgat^ neurons showed a positive correlation, whereas the vast majority, 36% (25 out of 70), had a negative correlation with sucrose’s palatability (Figures 3K and 3L; Positive vs. Negative chi-square test (1, 140) = 16.97, *p* < 0.0001). These results suggest that non-LHA^Vgat^ neurons (perhaps glutamatergic Vglut2 neurons) preferentially encode sucrose’s palatability with negative correlations (Figure 3M; Positive vs. Negative chi-square test (1, 135) = 11.46, *p* < 0.001).

### Activation of LHA^Vgat+^ neurons drives sucrose intake

If a subpopulation of LHA GABAergic neurons preferentially encodes sucrose’s palatability by increasing firing rates, its optogenetic stimulation should promote increased sucrose intake. To characterize its impact on sucrose intake, we used an open loop optogenetic stimulation involving 5 min blocks with laser and no-laser stimulation, while naïve VGAT-ChR2 mice had *ad lib* access to a sipper filled with sucrose 10% (Figure 4*A*). We analyzed the intake as a function of laser frequency (Figure 4B; two-way ANOVA, frequency by blocks interaction, F_(35, 96)_ = 2.268, *p* < 0.01), a *post-hoc* test uncovered that in the first three blocks with laser, 50 Hz stimulation induced a significant increase in sucrose intake (Figure 4B; Blocks 2, 4, and 6) compared with the previous blocks of no-laser (Figure 4*B*; *p* < 0.001; Blocks 1, 3, and 5). Figure 4*C* shows a raster plot for licking responses of a representative open loop session. At 50 Hz, we found that feeding (licking) was elicited within a 0.72 s ± 0.03 latency from laser onset, and it abruptly stopped (0.27 s ± 0.1) after laser offset (see blue dash rectangles). VGAT-ChR2 mice exhibited an increase in the laser-bound feeding as the laser frequency increased (Figure 4*D*; two-way ANOVA, group by frequency interaction, F_(5, 36)_ = 12.86, *p* < 0.0001). Thus, to induce sucrose intake, LHA^Vgat+^ neurons require continued activation. Our data then confirmed that 50 Hz is the best stimulation frequency to drive neuronal responses and sucrose intake.

**Figure 4.**
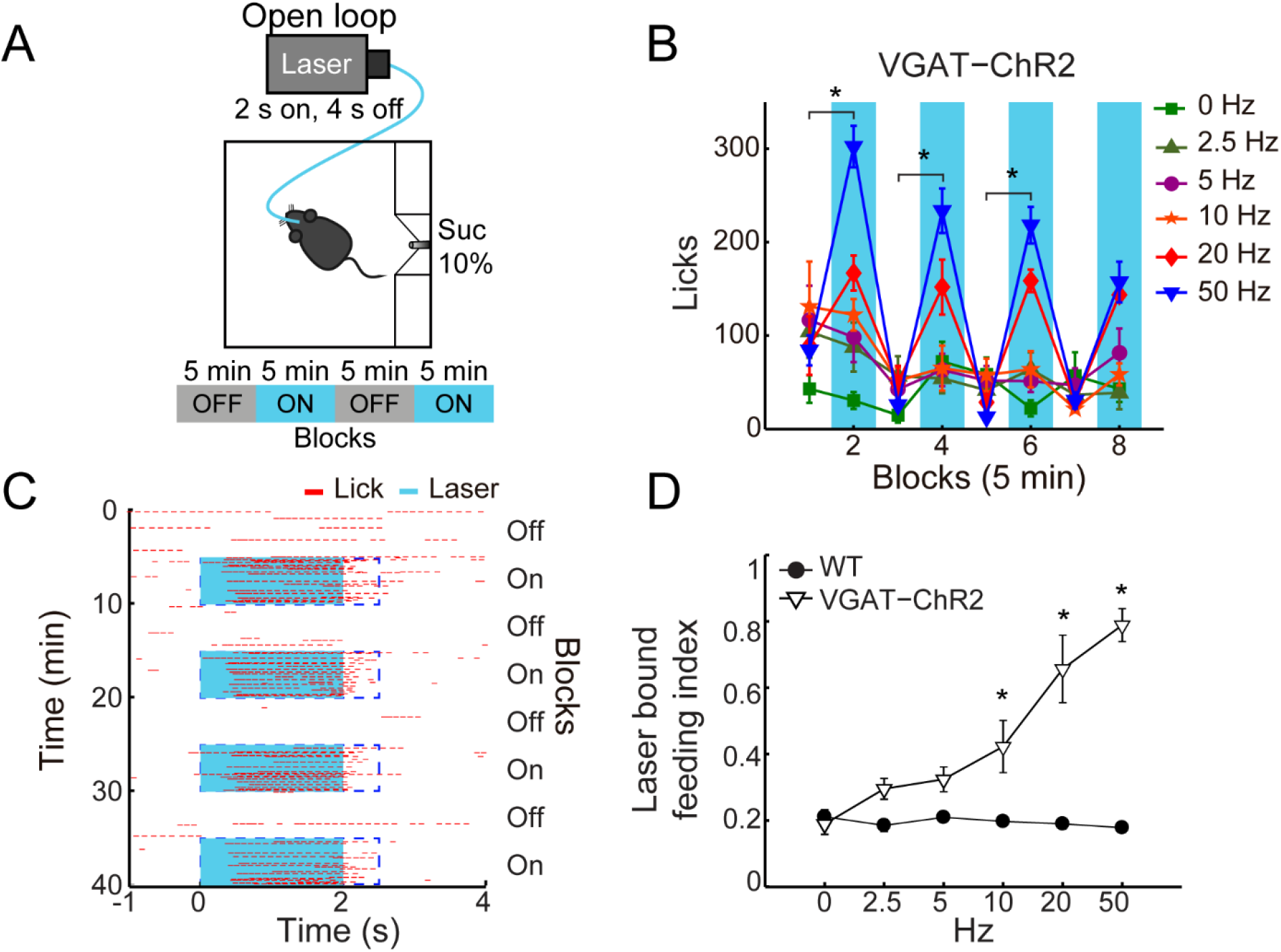
In sated VGAT-ChR2 mice, 50 Hz stimulation of LHA^Vgat+^ neurons induced the greatest intake of sucrose. ***A***, Behavioral setup for open-loop stimulation, where the laser was turned on regardless of the mice’s behavior. The box was equipped with a central sipper, where mice had free access to 10 % sucrose. Open-loop stimulation comprised 5 min block without laser followed by opto-stimulation for 5 min, in a cycle 2 s on – 4 s off, during a 40 min session. VGAT-ChR2 (n = 3) and WT (n = 5) mice received stimulation of only one of the following frequencies in ascending order per session 0, 2.5, 5, 10, 20, and 50 Hz. ***B,*** Opto-stimulation of LHA^Vgat+^ neurons induced sucrose (10%) intake in a frequency-dependent manner that peaked at 50 Hz. * Denotes statistically significant difference (*p* < 0.001) between non-laser and laser stimulated blocks at 50 Hz. Besides mice were sated, the lower intake at 0 Hz could be due to the lack of experience drinking in the novel box. ***C***, Representative raster plot of open-loop stimulation from a VGAT-ChR2 mouse at 50 Hz. Lick responses (red ticks) were aligned to laser onset (blue rectangles, opto-stimulation period). The laser bound index was the number of licks within 2.5 s from laser onset (dashed blue rectangles) divided by total licks. ***D***, The laser bound feeding index of VGAT-ChR2 increased as a function of the laser frequency. * Indicates significant difference from WT mice (*p* < 0.01). Using a Two-way ANOVA followed by the Holm Sidak test. In this and other figures, data represent mean ± SEM.

### LHA^Vgat+^ neurons do not induce a persistent hunger state

A previous study showed that brief optogenetic stimulation of “hunger-inducing” AgRP neurons in the arcuate nucleus before food availability promotes consummatory behavior that persists for several minutes in the absence of continued AgRP neuron activation (Burnett et al., 2016; Chen et al., 2016). Based on this study, we tested whether the pre-stimulation of LHA^Vgat+^ neurons could evoke a similar hunger state. Unlike AgRP neurons (Burnett et al., 2016; Chen et al., 2016), pre-stimulation of LHA^Vgat+^ neurons failed to evoke and sustain subsequent sucrose intake (see Supplementary Figure 4; two-way ANOVA, group by pre-stimulation protocol, F_(3, 56)_ = 0.50, *p* = 0.68). Our results suggest that feeding occurs while LHA^Vgat+^ neurons are active (Figure 4C), but they do not induce a persistent hunger state as AgRP neurons do (Chen et al., 2016).

### Open loop activation of LHA^Vgat+^ neurons drives intake of the highest sucrose concentration available

To test whether activation of LHA^Vgat+^ neurons promotes the intake of the most palatable available stimulus, naïve WT and VGAT-ChR2 mice were opto-stimulated in an open loop protocol, with access to three adjacent ports containing water, sucrose 3%, or sucrose 18% (Figure 5A). As expected, both WT and VGAT-ChR2 mice preferred sucrose 18% over water, and sucrose 3% (Figure 5B; WT: two-way ANOVA main effect tastants; F_(2, 1560)_ = 543.3, *p* < 0.0001; VGAT-ChR2: F_(2, 3144)_ = 983, *p* < 0.0001). However, we found that sated VGAT-ChR2 mice increased their intake of 18% sucrose mainly during the laser-activated blocks compared with controls (Figure 5B-*right*; two-way ANOVA group by laser block interaction F_(7, 1568)_ = 31.86, *p* < 0.0001) and rarely licked in the absence of stimulation (Figure 5C). In this regard, VGAT-ChR2 mice seem to counteract the evoked sucrose intake by voluntarily restraining consumption in no-laser blocks (Blocks 3, 5, and 7), resulting in no significant differences between groups in the total intake of 18% sucrose (Figure 5C, see black squares; unpaired Student’s t-test, *t* (196) = 1.512, *p* = 0.1320). A between days analysis revealed that the evoked 18% sucrose intake began from the first stimulation day (Figure 5D; RM ANOVA laser blocks, F_(1, 22)_ = 64.64, *p* < 0.0001), although sucrose consumption ramps up throughout the days (Figure 5D; RM ANOVA main effect days, F_(10, 220)_ = 2.169, *p* < 0.05). This suggests that LHA^Vgat+^ neurons, rather than inducing hunger *per se*, induced a learning process that potentiates the intake of the most palatable stimulus and confines consumption mainly in the presence of opto-stimulation.

**Figure 5.**
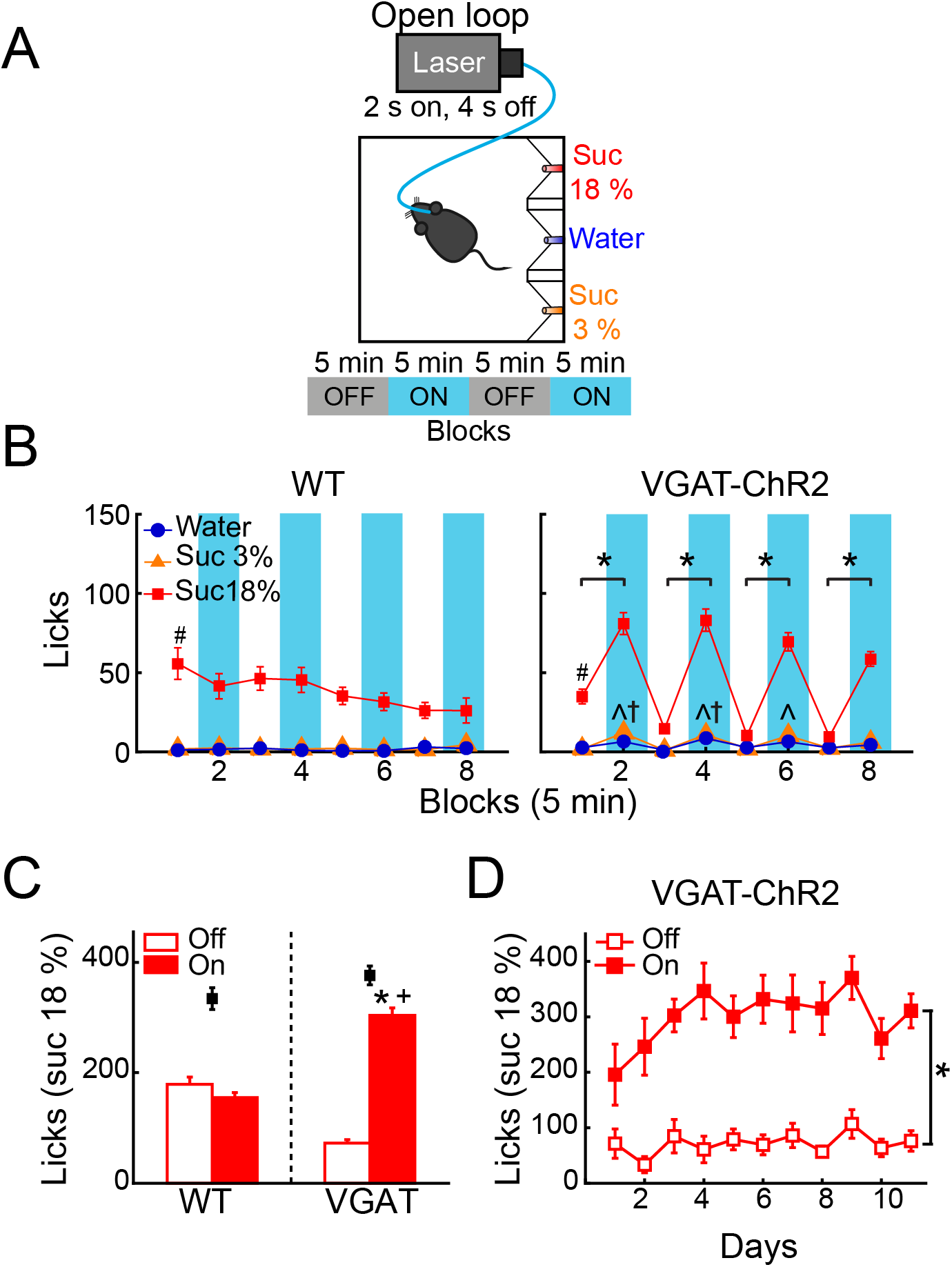
Open loop stimulation of LHA^Vgat+^ neurons biases consumption towards the most palatable tastant and reinforce intake during opto-stimulation. **(A)** Behavioral protocol. Sated mice (n = 6 WT and n = 12 VGAT-ChR2) were located in an operant box equipped with three ports. The left port contained sucrose 18%, the central port water, and the right port sucrose 3%, counterbalanced across individuals. For details of open loop, see Figure 4A. **(B)** *Left and right panel*, number of licks for each gustatory stimulus for WT and VGAT-ChR2 mice. Note that the VGAT-ChR2 mice increased consumption of the most palatable stimulus available (sucrose 18%) mainly in opto-stimulated blocks (blue rectangles), while it decreased intake in blocks without laser. **(C)** Mean licks for sucrose 18%, during blocks with laser on and off from WT and VGAT-ChR2 mice. Black squares show the total licks given for each group. **(D)** Licks for sucrose 18% during blocks with or without laser across days. * Indicates a significant difference (*p* < 0.0001) between laser and no-laser blocks. # *p* < 0.0001 statistically significant difference between sucrose 18% and the other stimuli. ^ *p* < 0.01 between WT and VGAT-ChR2 for sucrose 3%; † *p* < 0.01 higher intake of water of VGAT-ChR2 compared with WT;+ p < 0.0001 higher consumption of VGAT-ChR2 compared with WT for sucrose 18% during blocks with laser. Using a two-way ANOVA and RM ANOVA followed by the Holm Sidak test.

The small increase in water and sucrose 3% intake observed during opto-stimulation of LHA^Vgat+^ neurons could be explained by random stimulation near those tastants (Figure 5B *right panel*). Accordingly, we found in laser blocks 2, 4, and 6, a significant increase in water and sucrose 3% in the VGAT-ChR2 mice compared with the WT mice that completely neglected those tastants (Figure 5B, ^ *p* < 0.05 for sucrose 3%; † *p* < 0.05 for water). We hypothesize that this additional intake can be attributed to trials where the laser-activation occurred near these less palatable stimuli. To answer this, we employed a videography analysis (Figure 6A-B). When opto-stimulation occurred in the distance minor to 50 pixels (approximately 3 cm, see Materials and Methods) relative to sucrose 3% or water ports, the licking probability increased significantly after opto-stimulation (Figure 6B; Sucrose 3%: F_(9, 155)_ = 29.33, *p* < 0.0001; Water: one-way ANOVA; F_(9, 153)_ = 12.74, *p* < 0.0001). The angle of the head was less informative (Figure 6C-D; Sucrose 3%: F_(17, 147)_ = 0.71, *p* = 0.78; Water: one-way ANOVA; F_(17, 145)_ = 0.96, *p* = 0.49; Sucrose 18%: one-way ANOVA; F_(17, 153)_ = 0.49 *p* = 0.95). Conversely, since mice are naturally attracted to the most palatable tastant, they spent more time near the lateral port with 18% sucrose, increasing their overconsumption (Figure 6C, Video 1). Moreover, we observed that for the sucrose 18% port, even when the mouse position was twice as far (i.e., 6 cm) from sucrose 18% port, the licking probability increased significantly (Figure 6B *right panel*; one-way ANOVA, F_(9, 161)_ = 82.87, *p* < 0.0001). These data are consistent with a study showing that opto-stimulation of LHA^Vgat+^ neurons promotes the intake of the nearest stimulus (Nieh et al., 2016) but further demonstrates that the most palatable stimulus has nearly twice the distance of attraction than the other less palatable options.

**Figure 6.**
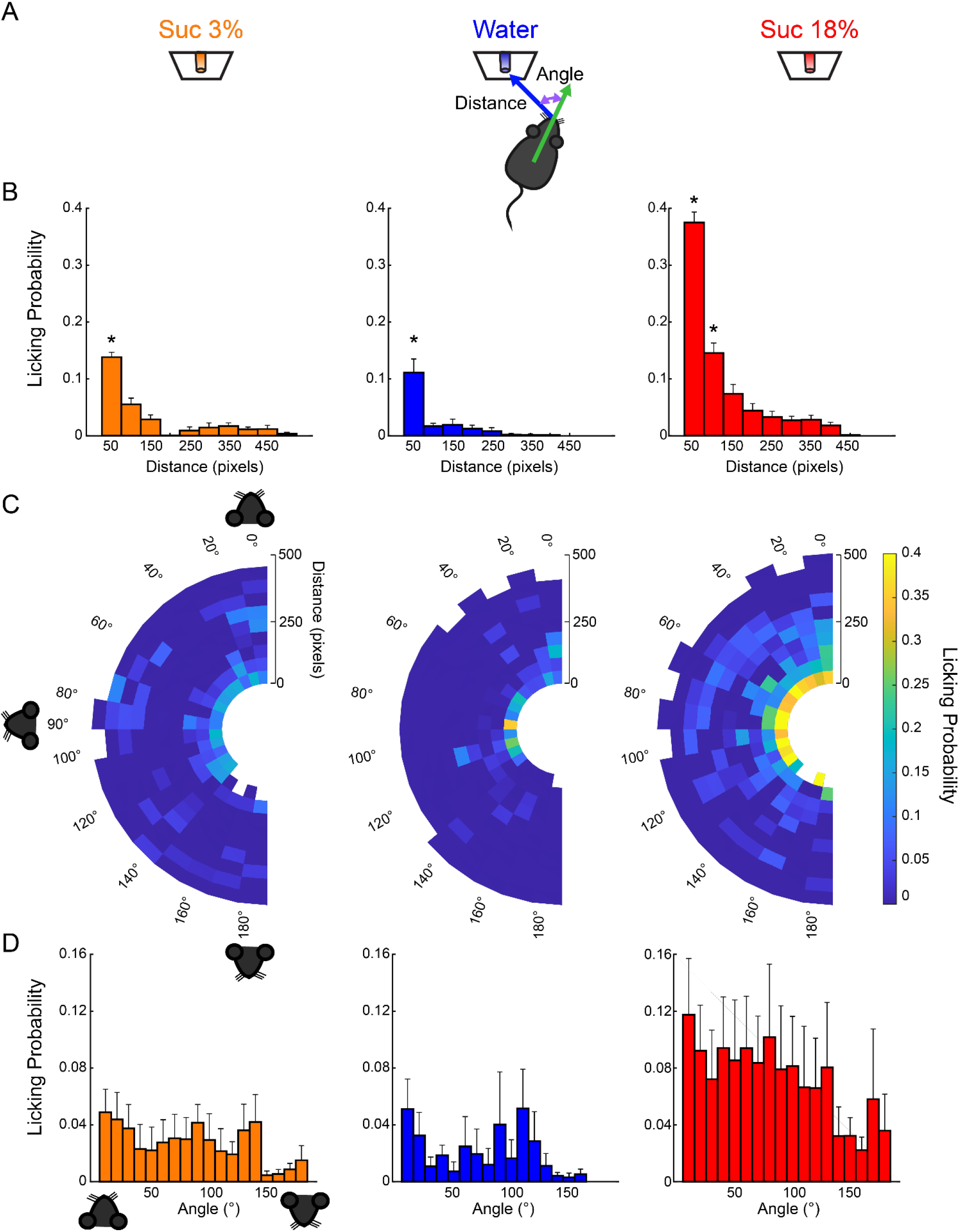
Licking probability increased as a function of distance. **(A)** Schematics for video tracking (see Materials and Methods). We calculated the distance and the head angle between the mouse’s nose and each of the three sippers containing sucrose 3%, water, or sucrose 18% for each laser onset. **(B )** It depicts the probability that laser onset evoked licking as a function of the mouse’s distance relative to each sipper. **(C)** Polar plots depict the licking probability after laser onset, given the distance and angle of the mouse relative to a licking sipper. At the onset of each laser stimulation, we computed the head angle and distance relative to each sipper, and, in the following 6 s window, we tally if the mouse licked to any sipper. We normalized by dividing the number of times licking was elicited by the number of first lasers occurring on those coordinates. The drawing of the mice’s head illustrates that if a mouse was facing towards the sipper, the angle is 0°, in a perpendicular position (90°), or 180° if it was looking in the opposite direction. **(D)** The probability of licking after laser onset versus the angle between head direction and the sipper. * Indicates significantly different (p < 0.001) from all other distances. One-way ANOVA followed by the Holm Sidak test.

### Open loop activation of LHA^Vgat+^ neurons also increases the time spent and drives the intake of the most palatable solid food available

We also explored whether LHA GABA neurons could induce the intake of the most palatable solid food available. Thus, using an open loop protocol, we opto-stimulate these neurons while mice choose among different solid foods. We found that optogenetic activation of LHA^Vgat+^ neurons increased the time spent near the most palatable food available. VGAT-ChR2 mice spent more time near the high-fat diet than the granulated sugar cube or the Chow food pellet, relative to WT group. Also, the intake of transgenic mice was higher for high-fat diet than the other food stimuli (see Supplementary Figure 5 and Video 2; Time spent: one-way ANOVA, F_(5, 306)_ = 38.52, *p* < 0.0001; Intake: one-way ANOVA, F_(5, 282)_ = 68.76, *p* < 0.0001), however, both WT and VGAT-ChR2 mice consumed similar amounts of high-fat diet (*p* = 0.0985.), perhaps because it is a highly palatable food. When mice were able to choose between a sugar cube and a Chow pellet, now the activation of these neurons increased the time spent and consumption of sugar cube over Chow (see Supplementary Figure 6 and Video 3; Time spent: one-way ANOVA, F_(3, 96)_ = 16.13, *p* < 0.0001; Intake: one-way ANOVA, F_(3, 92)_ = 13.65, *p* < 0.0001). Our data suggest that open loop stimulation of LHA GABAergic neurons promotes the attraction to and the intake of the most palatable food available.

### Closed-loop stimulation of LHA^Vgat+^ neurons drives the intake of the nearest appetitive stimuli

To further test the idea that LHA^Vgat+^ neurons could induce a preference for a proximal (but less palatable) stimulus over a distal one with higher hedonic value, we used a closed-loop stimulation protocol. That is, the same mice (from Figure 5) were photostimulated, but here the laser was triggered by each head entry in the central port (i.e., opto-self-stimulation). This closed-loop configuration guarantees that stimulation only occurs proximal to water, the least palatable stimulus of the three in the box (Figure 7*A-right*). We found that activation of LHA^Vgat+^ neurons is rewarding since transgenic mice visited the central port significantly more and performed a higher number of opto-self-stimulations than the WT mice (Figure 7B; unpaired Student’s t-test, *t*(196) = 12, *p* < 0.0001). Surprisingly, the total number of licks in the session seen in Figure 7C shows that during closed-loop stimulation, the VGAT-ChR2 mice explored and consumed more water than the WT mice (two-way ANOVA group by tastants interaction, F_(2, 585)_ = 13.09, *p* < 0.0001, *post hoc* test at water, p < 0.0001). Note that in both open loop and closed-loop protocols, the WT and VGAT-ChR2 mice exhibited, at the end of the session, a similar number of licks for 18 wt% sucrose (Figure 7C; two-way ANOVA group by protocol interaction F_(1, 391)_ = 2.172, *p* = 0.1414). That is, the total licks for sucrose 18% were not significantly different between groups, suggesting that these neurons do not interfere with the overall attractiveness of sucrose. However, by counting the number of licks given when the laser was turned on (in a 2.5 s window from laser onset- see Figure 7D), we uncovered a striking change in preference. As noted, in the open loop, the VGAT-ChR2 mice exhibited a higher preference for sucrose 18 wt%, over both sucrose 3 wt% and water. Now, in the closed-loop protocol, the same transgenic mice consumed more water compared with WT (two-way ANOVA group by tastants interaction, F_(2, 585)_ = 12.18, *p* < 0.0001, post hoc test at water, *p* < 0.0001), and the consumption of water was about the same as sucrose 18% (*p* = 0.8731, n.s.; Figure 7D). The increase in water intake was selective to the photostimulation window since the transgenic mice only drank sucrose 18% and neglected water when the laser was turned off (see arrow Figure 7E; two-way ANOVA group by tastants interaction, F_(2, 585)_ = 64.16, *p* < 0.0001, post hoc test at water vs. sucrose 18%, p < 0.0001). Thus, LHA^Vgat+^ neurons could promote water intake, but only when it is the nearest stimulus. The sucrose preference index (Figure 7F) showed that during the open loop, all VGAT-ChR2 mice preferred sucrose 18% over water (Figure 7F; values > 0.5 and near to 1), whereas in the closed-loop protocol, when water was the nearest stimulus, most mice (n = 9 out of 12), significantly diminished their sucrose preference (Figure 7F, see the drop in preference index; paired Student’s *t*-test, *t* (58) = 7.98, *p* < 0.0001). From these 9 animals, 6 exhibited a higher preference for water over sucrose 18% (Preference index values < 0.5; Video 4). Figure 7G shows the color-coded location of fiber optic tips in each VGAT-ChR2 mice plotted in Figure “7F.” However, the location of the optical fibers does not explain the variability in preference. Perhaps the variability is due to different behavioral strategies used by each mouse. Nevertheless, the large variability in the drop of sucrose preference resembles the findings with sweet-induced facilitation by electrical ICSs (Poschel, 1968). These data suggest that LHA^Vgat+^ neurons drive consummatory behavior by integrating the stimulus’s proximity and hedonic value.

**Figure 7.**
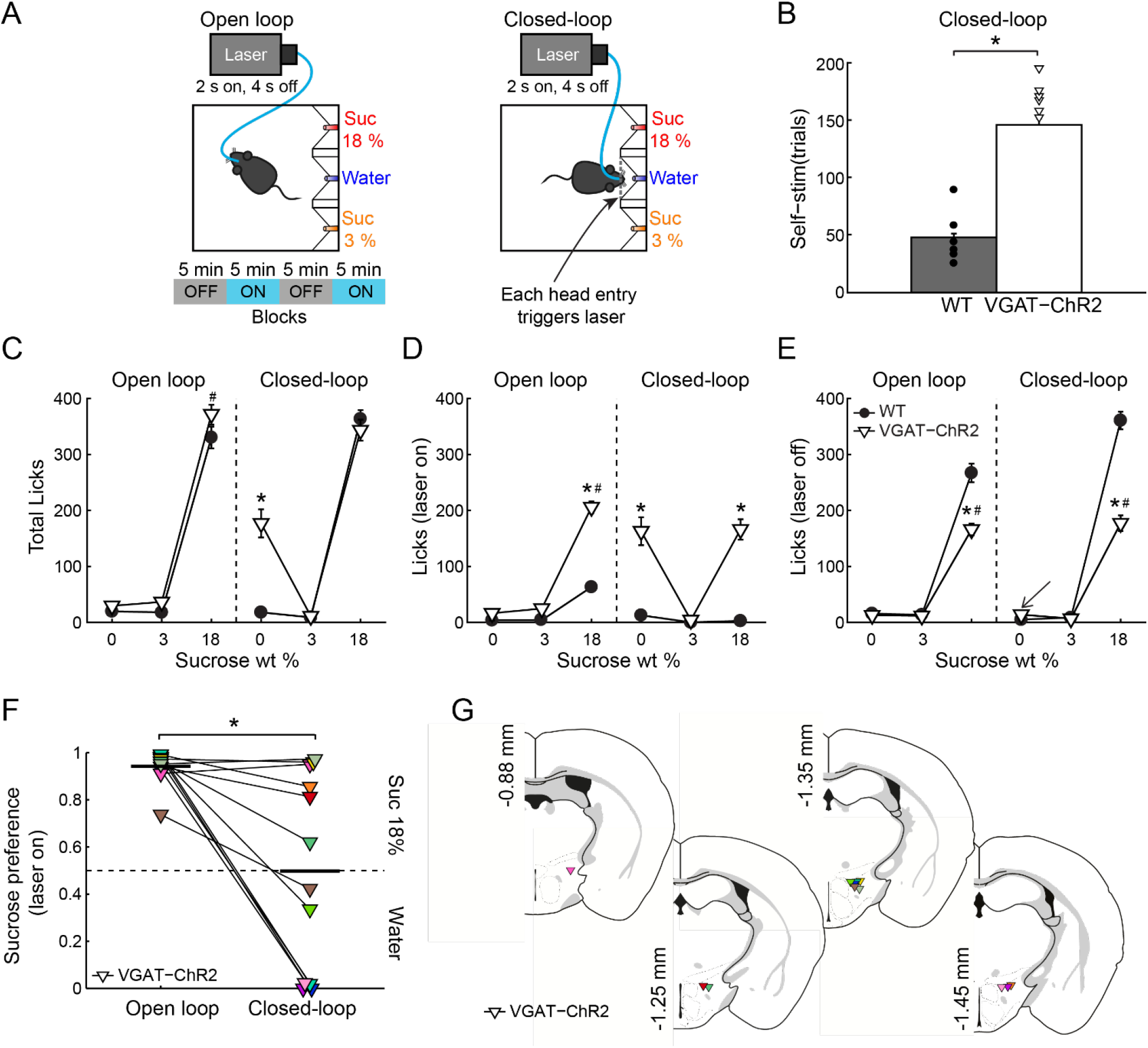
If stimulation occurred near the less palatable stimulus, LHA^Vgat+^ neurons induced water intake despite sucrose. **(A)** For the open loop stimulation, the laser was activated (2s on, 4s off) regardless of behavior and mice position. Same mice and conventions as in Figure 5*A*. In the closed-loop configuration, the laser was triggered by a head entry in the central port. **(B)** The number of laser activations during the closed-loop protocol. The VGAT-ChR2 mice self-stimulate more than the WT mice. Each dot and triangle represent a single mouse. Unpaired t-test. **(C)** The total number of licks given for each stimulus during the entire session. In the open loop both mice groups licked more sucrose 18% than water or sucrose 3%. However, in the closed-loop configuration, the VGAT-ChR2 consumed more water (0 % sucrose) than the WT mice. **(D)** The number of licks evoked during 2.5 s after laser onset for each gustatory stimulus. **(E)** The number of licks when laser was turned off for each gustatory stimulus. **(F)** The water-sucrose 18% preference index from VGAT-ChR2 mice during photostimulation was defined as the number of 18% sucrose licks divided by total licks for sucrose 18% + water. Thus, values higher than 0.5 indicate sucrose 18%, and values lower than 0.5 prefer water. Some VGAT-ChR2 mice preferred water over sucrose. The horizontal black lines indicate mean preference in both stimulation protocols. Paired t-test. **(G)** Fiber optic location in the LHA of VGAT-ChR2 mice. * *p* < 0.0001 indicates a significant difference from WT mice and stimulation protocols. # *p* < 0.0001 between sucrose 18% and the other stimuli. Two-way ANOVA followed by the Holm Sidak test.

### After repeated stimulation, the correlation between laser-bound feeding and self-stimulation strengthens

Previous studies have shown that after repeated LHA bulk stimulation (either electrically or with optogenetic targeting all cell-types together), subjects switched from exhibiting stimulus-bound feeding to only self-stimulating, indicating that LHA evoked feeding and reward could represent two distinct processes (Gigante et al., 2016; Urstadt and Berridge, 2020). Consequently, we explored whether laser-bound feeding changed after repeated optogenetic stimulation. In contrast to previous studies, we observed that laser-bound feeding induced by LHA^Vgat+^ neurons strengthened across sessions (Figure 8A, First 3 days: r = 0.12, p = 0.69; Last 5 stimulation days: r = 0.63, p < 0.05) and after repeated stimulation, both opto-self-stimulation and laser-bound feeding exhibited a robust correlation (Figure 8B, First 3 days: r = 0.42, p = 0.16; Last 5 days: r = 0.61, p < 0.05), suggesting that LHA evoked-feeding (licking) involves a learning process (Sharpe et al., 2017) and that LHA GABAergic neurons are a common neural substrate for feeding and reward.

**Figure 8.**
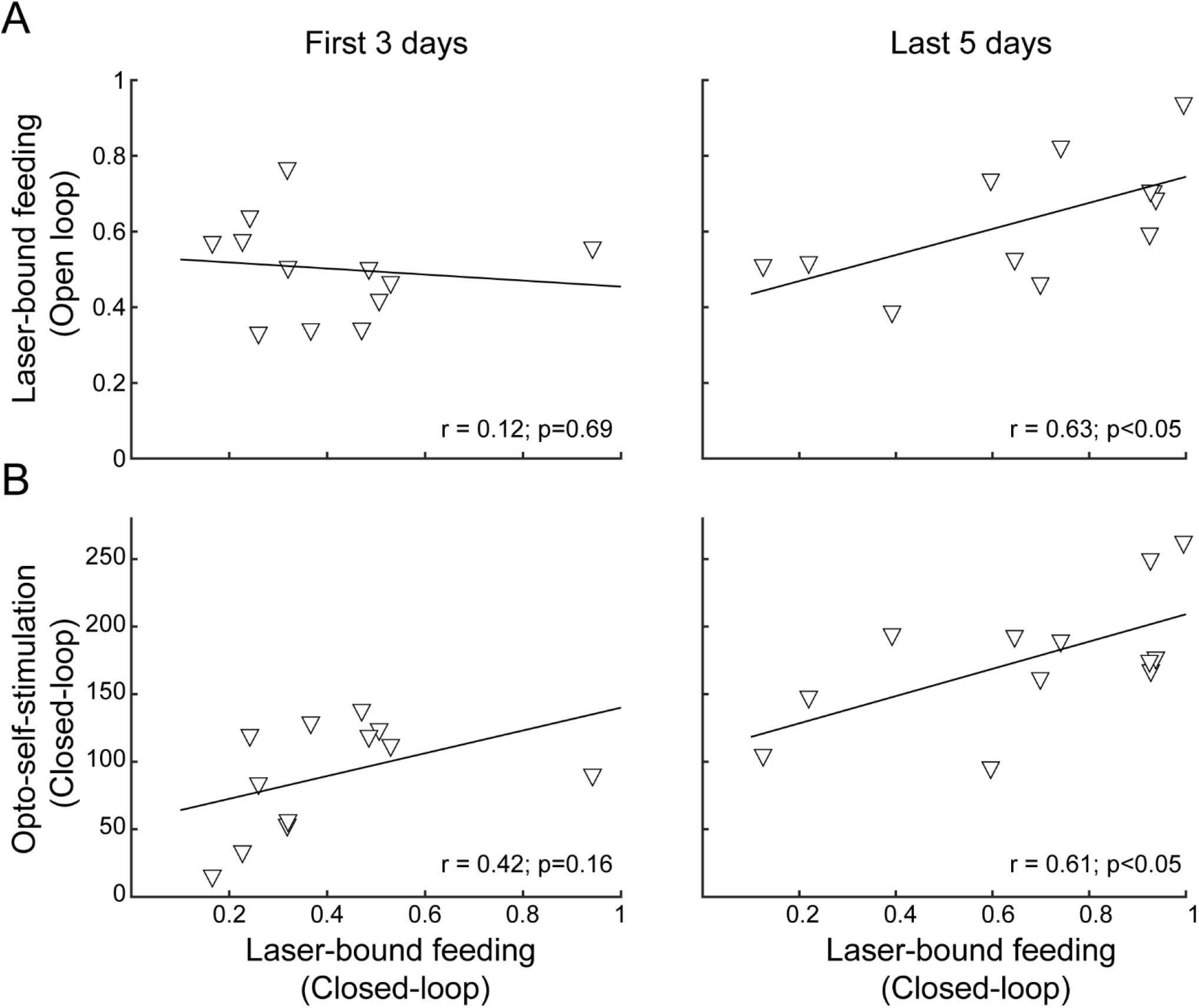
After repeated activation of LHA^Vgat+^ neurons, laser-bound feeding (in open vs. closed-loop) strengthened and exhibited a positive correlation. **(A)** Scatter plots showing the relationship between laser-bound feeding (licks given during the 2.5 s window from laser onset relative to total licks in the session) in both open loop and closed-loop tasks during the first 3 days (l*eft panel*) and the last 5 days of opto-stimulation of open loop vs. opto-self-stimulation of the closed-loop protocols (*right panel*). **(B)** Likewise, after repeated activation, both opto-self-stimulations and laser-bound feeding (licking) showed a significant correlation. Scatter depicting the average number of opto-self-stimulations vs. the laser-bound feeding observed in the closed-loop protocol.

### The “stimulus-proximity effect” is not restricted to liquid tastants, and it also occurs with chocolate and Chow pellets

The “proximity effect” evoked by LHA^Vgat+^ neurons was not restricted to liquid tastants, it also applied to solid foods. In a real-time place preference arena with four stimuli (Figure 9A), when chocolate pellets or Chow food were the designated foods near opto-self-stimulation of LHA^Vgat+^ neurons, mice also increased the time spent near those foods (Figure 9B; Pellet: one-way ANOVA; F_(4, 145)_ = 99.05, *p* < 0.0001; Chow, F_(4, 145)_ = 182, *p* < 0.0001) as well as their intake (Figure 9C; Pellet: unpaired Student’s t-test, *t*(68) = 3.234, *p* < 0.05; Chow: unpaired Student’s t-test, *t*(68) = 3.651, p < 0.05), see Video 5. Thus, these LHA GABAergic neurons reinforced the approach and exploration to any, if not the most appetitive stimulus, that happened to be proximal to the opto-stimulation.

**Figure 9.**
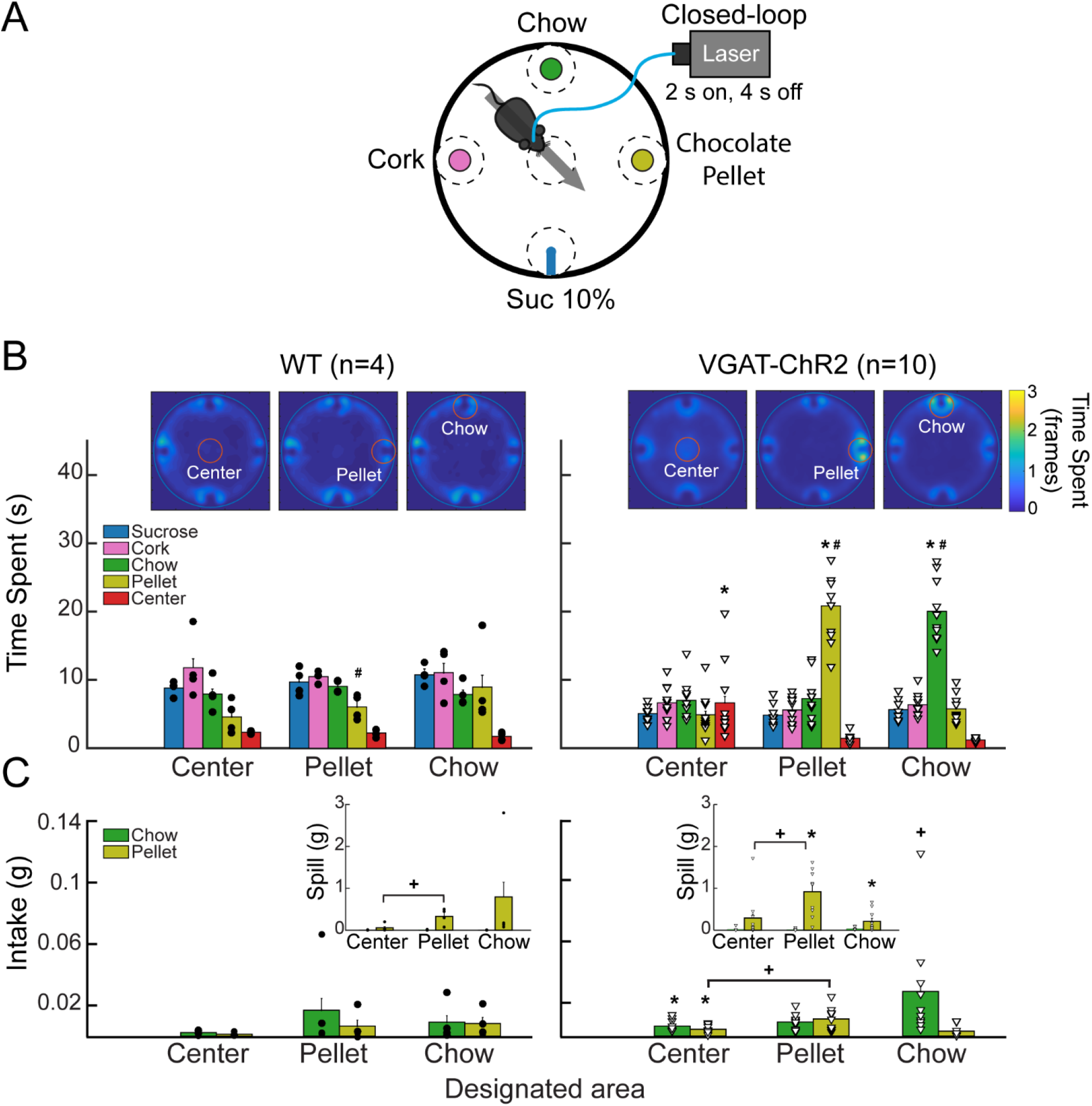
Stimulation of LHA^Vgat+^ neurons is rewarding and biases approach towards the nearest stimulus. **(A)** Different stimuli (plates containing cork, Chow, chocolate pellets, or a sipper filled with 10% liquid sucrose) were presented simultaneously in a circular arena. In this task, when mice cross a designated area (dashed circles), the laser was turned on (2 s on; 4 s off) in a 40 min session. Thus, a mouse had to leave and re-enter the designated area to receive a new opto-self-stimulation. Only one designated area was used per session, and it remained in the same position for up to 3 or 4 consecutive sessions. **(B)** Heat map and time spent from a representative WT (left panel) and VGAT-ChR2 mice (right panels), when the designated area was either the center, the pellet, or the Chow (see red circles for the currently designated zone). Colorbar indicates the number of frames the subject was detected in a given pixel; higher values indicate it remained in the same place for a longer time. *Below*, bar graphs depict the time spent in seconds exploring the designated area (within a radius of 5 cm). Each dot and triangle represent a single individual. Though mice normally avoid exploring the center of an open field, opto-self-stimulation in the center zone increased the time transgenic mice spent exploring it, compared with WT. Transgenic mice spent more time exploring the chocolate pellets and the Chow food when they were in the designated zones. **(C)** Intake of chocolate pellet and Chow. Activation of LHA^Vgat+^ neurons in the center zone increased chocolate pellets and chow consumption compared with WT. *Inset*: Spill from chocolate pellets and chow. * Indicates statistically significant difference (*p* < 0.05) from WT. + p < 0.05 shows a significant increase in intake and spill during the session that a designated area was opto-self-stimulated relative to the sessions where the center was opto-self-stimulated. Unpaired Student *t*-test. # *p* < 0.01 indicates a significant difference between designated area opto-stimulated and the other open field areas. One-way ANOVA followed by the Holm Sidak test. See Video 5.

### LHA^Vgat+^ neurons activation is rewarding

In the same WT and VGAT-ChR2 mice tested in the closed-loop configuration seen in Figure 7, we went on to show that mice visited the central water-port to opto-self-stimulate. To do this, we performed extinction sessions with the laser disconnected (Figure 10A). We observed a rapid decrease in the number of opto-self-stimulations (Figure 10B; Extinction phase, gray shadow). As expected, self-stimulation rapidly recovered when the laser was turned on again (Figure 10, see water after extinction). Thus, LHA^Vgat+^ neurons convey a hedonically positive, rewarding signal (Jennings et al., 2015).

**Figure 10.**
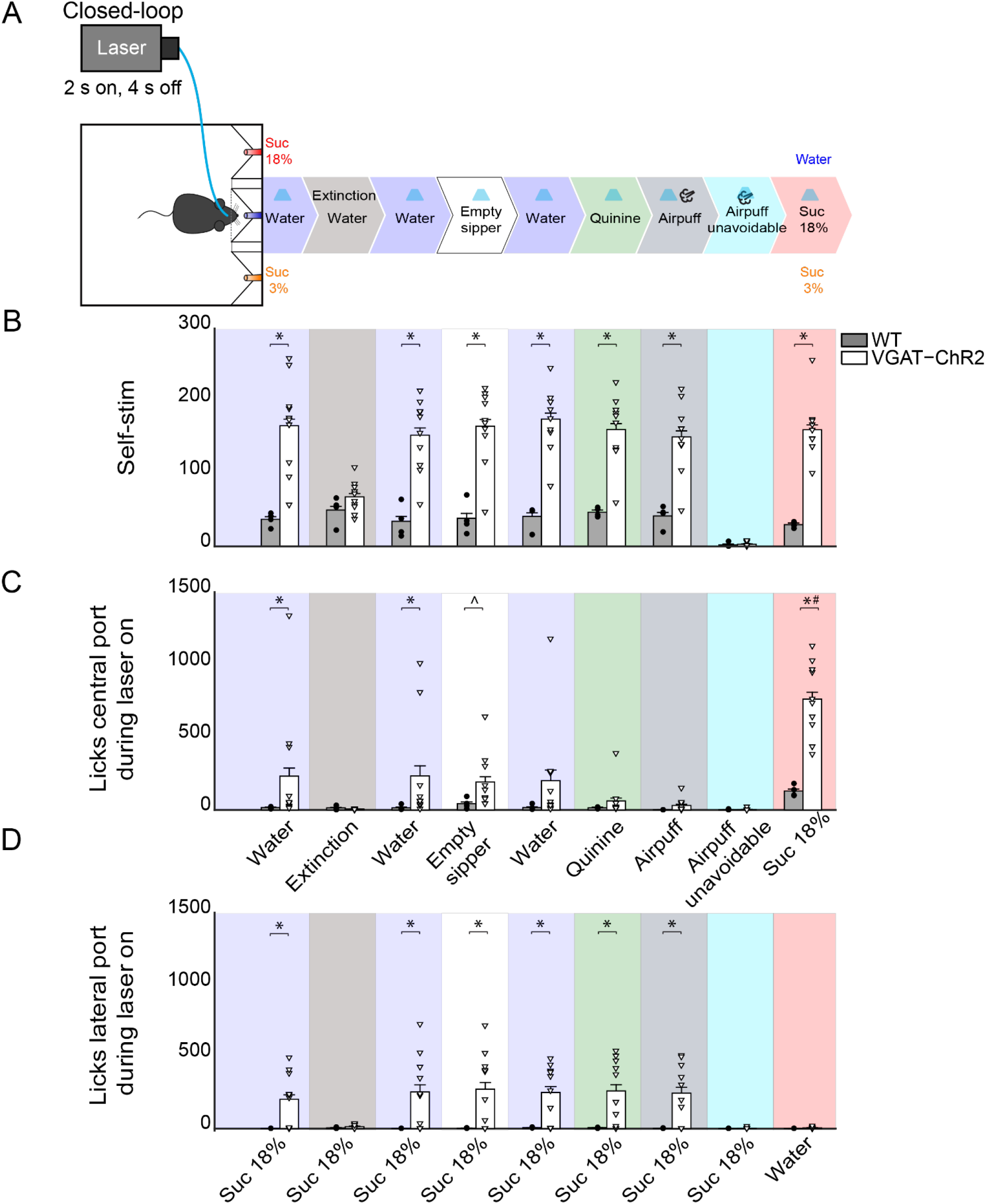
Activating LHA^Vgat+^ neurons does not induce the intake of a bitter tastant, nor an unavoidable aversive stimulus, but it increases sucrose consumption. **(A)** Schematic of behavioral setup showing stimuli delivered at central port. The water stimulus in the central port was replaced by an empty sipper, quinine, or airpuff. In all phases, head-entry in the central port triggered the laser, except in the airpuff unavoidable condition, where the first lick delivered both the laser and the airpuff. Finally, the airpuff in the central port was replaced by sucrose 18%. **(B)** The number of opto-self-stimulations given for each stimulus. The VGAT-ChR2 mice performed more self-stimulations than WT, except during extinction sessions and when the airpuff was unavoidable. **(C)** The number of licks given to the central port. The licks in the central port decreased when an aversive stimulus was present, such as quinine or airpuffs. In contrast, a non-edibles stimulus such as an empty sipper elicited more licks from VGAT-ChR2 mice (*p* < 0.05). Moreover, when sucrose 18% was in the central port, VGAT-ChR2 group increased its consumption substantially compared with WT. **(D)** The number of licks in the lateral port containing sucrose 18% during opto-self-stimulation. The intake of the lateral port of sucrose 3 % is not shown because it was neglectable. Each dot and triangle represent a single individual. ^ Denotes statistically significant difference (*p* < 0.0001) from WT. Unpaired-Student t-test. * *p* < 0.05 relative to WT group. # *p* < 0.05 between sucrose 18% from other stimuli delivered at the central port. Two-way ANOVA followed by the Holm Sidak test.

### LHA^Vgat+^ neurons also promote licking an empty sipper

A previous study demonstrated that chemogenetic activation of LHA^Vgat+^ neurons increased gnawing to non-edible cork (Navarro et al., 2016). In the absence of other stimuli but cork, we also observed gnawing behavior in some mice, although we did not systematically study stereotypy (see Video 6). Instead, we further explored this idea by replacing water, in the central port, with an empty sipper (Figure 10, Empty sipper). Transgenic mice continued opto-self-stimulating at the same place as if water were still present in the central port. LHA^Vgat+^ stimulation evoked licking an empty sipper compared with WT mice (Figure 10C, unpaired Student’s t-test, t (40) =2.532, *p* < 0.05). Thus, LHA^Vgat+^ neurons evoke an imminent urge to express consummatory behavior even in the form of dry licking an empty sipper, a non-biological relevant stimulus, but only when it is the nearest stimulus to photostimulation. This effect is perhaps also mediated by an increase in the rewarding value of appetitive oromotor responses *per se*.

### When sucrose is available, activation of LHA^Vgat+^ neurons neither promotes liquid intake of an aversive bitter tastant nor tolerance of punishment, but it further increases sucrose consumption

To explore whether the “stimulus-proximity effect” was also applied to aversive stimuli, we replaced the central stimulus (water) with aversive stimuli, including quinine (a tastant that humans experience as bitter taste) or airpuffs (Figure 10*A*). Upon quinine presentation, the number of licks given in the central port sharply decreased (Figure 10C, green), but transgenic mice continued to self-stimulate and go to the lateral port to lick for sucrose 18% (Figure 10D, green). We obtained similar results to those in quinine stimulation during the airpuff delivery phase (Figure 10D, dark gray). Next, we required the animal to lick, at least once, the central sipper to trigger opto-stimulation (thus making the airpuffs unavoidable). Under this condition, transgenic mice completely stopped self-stimulation (Figure 10B, cyan) and aborted 18% sucrose intake from the lateral port (Figure 10D, cyan). These results suggest that, in sated mice, LHA^Vgat+^ neurons did not induce quinine intake nor increase tolerance to airpuffs.

Finally, we explored whether proximity to the most palatable tastant further facilitated its overconsumption. Thus, we exchanged the position of water and sucrose 18%. When sucrose 18% was delivered in the central port, transgenic mice greatly overconsumed it (Figure 10C, pink). The intake of sucrose 18% was higher than for all the other tastants tested previously (*p* < 0.0001). These results collectively suggest that activation of LHA^Vgat+^ neurons is rewarding and promotes increased consumption of the nearest stimulus even if it is not the most palatable (e.g., water and empty sipper). If the nearest stimulus happens to be the most palatable (i.e., sucrose 18%), then LHA GABA neurons further facilitated its consumption.

### LHA GABA neurons increased quinine intake but only in water-deprived mice

Having demonstrated that in the presence of sucrose, activation of these neurons failed to increase quinine intake (Figure 10C green), with a new group of naïve mice, we then explored whether LHA^Vgat+^ neurons could induce quinine intake when it was the only option. As expected, in sated mice, we found that activation of LHA^Vgat+^ neurons did not affect quinine intake compared with WT (Figure 11A; unpaired Student’s t-test, t (46) =0.9925, p = 0.3262). Surprisingly, it promoted a higher quinine intake when transgenic mice were water-deprived (Figure 11B; unpaired Student’s t-test, t (46) =2.958, p < 0.01), specifically during photostimulation window (Figure 11B; unpaired Student’s t-test, t (46) = 4.473, p < 0.0001), suggesting that activation of these neurons is sufficient to increase the acceptance of bitter tastants but only during water deprivation. In contrast, we found that regardless of homeostatic needs, activation of these neurons increased sucrose 18% intake relative to WT mice (Figure 11C-D; unpaired Student’s t-test, t (30) = 6.933, p < 0.0001; unpaired Student’s t-test, t (30) =3.183, p < 0.01).

**Figure 11.**
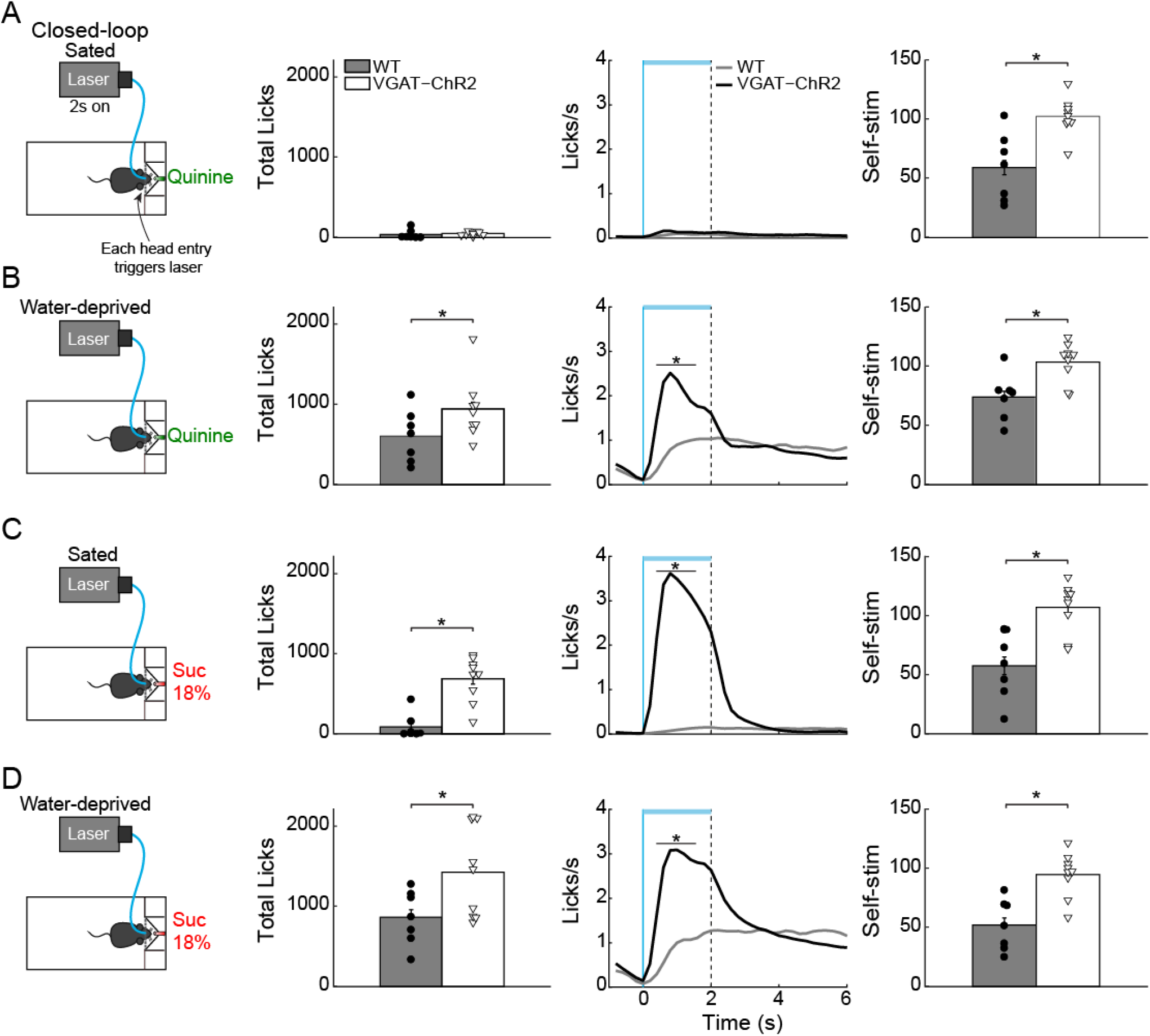
Water deprivation gates a time window where activation of LHA GABAergic neurons increases quinine intake. **(A)** Sated mice had free access to one sipper filled with quinine. The laser was turned “on” in closed-loop protocol (*left panel*). Each head entry triggers 2 s laser “on” followed by a 4 s time out with no-laser. In sated VGAT-ChR2 mice, activation of these neurons did not increase quinine intake (*central panel* shows the PSTH of the lick rates aligned to laser onset, time = 0s). Although sated transgenic mice do not lick for quinine, they continued opto-self-stimulating (*right panel*). **(B)** Water deprived transgenic mice consumed more quinine than WT mice (*central panel*) when the laser was turned on (horizontal blue line). **(C-D)** Total licks, PSTH of the lick rate (*central panels*), and number of opto-self-stimulations of 18% sucrose during sated and water deprivation condition (*right panel*). The horizontal blue line indicates opto-self-stimulation window. The vertical blue line indicates the laser onset, and the black dashed line the laser offset. * *p* < 0.01 indicates significant differences compared with WT mice according to an unpaired t-student test.

### Activation of LHA^Vgat+^ neurons enhances palatability

We next explored whether, at equal stimulus distance, these neurons could enhance palatability responses. To do this, in a new group of water-deprived mice, we employed a brief access test again. In this task, we delivered gustatory stimuli water, sucrose 3%, and sucrose 18% from the same sipper tube, but in different trials. Figure 12*A* displays the trial’s structure (same conventions as in Figure 2). We found that both groups increased their licking rate as a function of sucrose’s concentration, reflecting its palatability (Figure 12B; two-way ANOVA main effect of tastants, F_(2, 174)_ = 9.101, *p* < 0.001). However, during the 2 s of laser stimulation, the VGAT-ChR2 mice exhibited a greater lick rate for all three tastants. But, when the laser was turned “off,” these mice abruptly stopped licking for water and sucrose 3% compared with the WT group (Figure 12*B*, *right* panel; two-way ANOVA group by tastants interaction, F_(2, 174)_ = 5.293 *p* < 0.01). Moreover, when stimulation was paired with water trials, transgenic mice selectively increased their lick rate to water relative to WT (Figure 12C, blue line, and arrow; two-way ANOVA group by tastants interaction, F_(2, 84)_ = 6.009, *p* < 0.01), even surpassing licking responses evoked by the most palatable sucrose 18%. A similar enhancement of oromotor responses was observed by pairing sucrose 3% trials with LHA^Vgat+^ opto-self-stimulation (Figure 12D, orange; two-way ANOVA group by tastants interaction, F_(2, 84)_ = 16.72, *p* < 0.0001). Likewise, the laser-bound feeding (licking rate) was strongest when sucrose 18% was paired with the optogenetic stimulation (Figure 12E, red, see arrow; two-way ANOVA group by tastants interaction, F_(2, 84)_ = 16.49, *p* < 0.0001).

**Figure 12.**
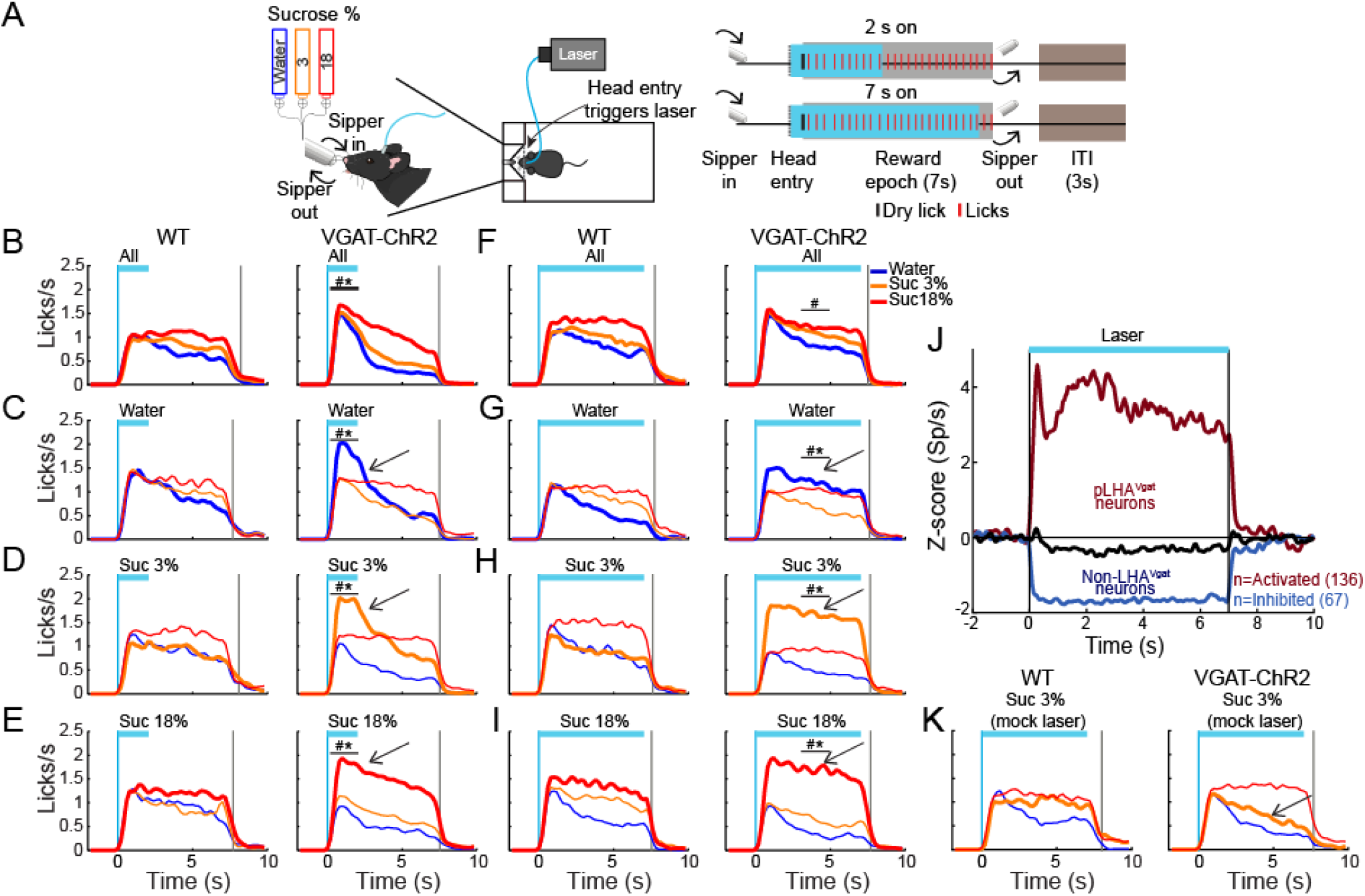
Activation of LHA GABA neurons enhances water and sucrose’s palatability. **(A)** Closed-loop stimulation during the brief access test. *Left panel,* behavioral setup. The box contained a sipper that delivered a ~2 μL drop of either water, sucrose 3%, or 18% per lick. *Right panel*, schematics of the structure of a trial. A head entry in the port (dashed vertical line) triggered opto-self-stimulation, at 50 Hz, in some sessions for 2 s and others for 7 s (blue rectangles). The reward epoch begins with the first dry lick on the sipper (black tick) and always last 7s (gray rectangles). During the reward epoch, a drop of tastant was delivered in each lick (red marks). At the end of the reward epoch, the sipper was retracted, and an Inter Trial Interval (ITI) of 3 s began. **(B-E)** Data of closed-loop for 2s stimulation in the Reward epoch. PSTH of lick rate of WT (left panel) and VGAT-ChR2 mice (*right panel*) aligned (Time = 0 s) to laser onset. Opto-self-stimulation was delivered during all trials (B) or during only water trials (C, see arrow), or sucrose 3% (D), or 18 % (E). **(F-I)** Licking responses but during 7 s opto-self-stimulation. Same conventions as in B-E. The vertical blue line indicates laser onset and gray line the end of the reward epoch. The horizontal blue line indicates opto-self-stimulation window. * Indicates significant difference (*p* < 0.05) from WT group. # *p* < 0.05 among stimuli. Two-way ANOVA followed by the Holm Sidak test. ***J,*** Normalized Z-score population activity of 284 neurons recorded in LHA in VGAT-ChR2 mice (n = 3). The laser was turned on for 7 s at 50 Hz. Neuronal responses were aligned to laser onset (Time = 0 s). Red and blue colors indicate activated or inhibited laser-evoked responses, relative to baseline (−1 to 0 s). Black trace illustrates neurons unmodulated by the laser. **(K)** Mock laser stimulation did not affect palatability responses. PSTH of licking responses, opto-self-stimulation was delivered along with sucrose 3% trials during 7s (arrow). Note that both groups increased the lick rate as a function of palatability, same conventions as in B-E. To elicit mock laser stimulation, mice were connected to a mock fiber optic (with no-laser connected), whereas the real fiber was glued outside the mice’s head to emit blue light.

Finally, we show that palatability responses could be artificially extended as long as LHA^Vgat+^ neurons were continuously activated. For this, we photostimulated them for up to 7 s. We observed a similar enhancement pattern, maintained for the duration of opto-stimulation (see Figure 12F-I see arrows; All trials: two-way ANOVA main effect of tastants, F_(2, 114)_ = 12.53, *p* < 0.0001; Water trials: two-way ANOVA group by tastants interaction, F_(2, 84)_ = 9.850, *p* < 0.001; Sucrose 3% trials: two-way ANOVA group by tastants interaction, F_(2, 84)_ = 33.47, *p* < 0.0001; Sucrose 18% trials: two-way ANOVA group by tastants interaction, F_(2, 84)_ = 8.360, *p* < 0.001), suggesting that LHA^Vgat+^ neurons can adjust the enhancement of oromotor palatability responses by simply sustaining its neuronal activity. We verified that using optrode recordings, 7 s optogenetic stimulation produced sustained LHA^Vgat+^ neuronal responses (Figure 12J). Moreover, transgenic mice did not merely use the light as a cue to guide behavior since laser-stimulation with a mock optical fiber failed to increase licking (Figure 12K; two-way ANOVA group by tastants interaction, F_(2, 54)_ = 1.619, *p* = 0.207). In sum, our data demonstrate that activation of LHA^Vgat+^ neurons triggers a reinforcing signal that amplifies the positive hedonic value of proximal stimuli, promoting overconsumption.

## Discussion

LHA has historically been viewed as a critical center for feeding (Anand and Brobeck, 1951; Delgado and Anand, 1953; Teitelbaum and Epstein, 1962), although it also processes sucrose’s palatability related information (Norgren, 1970; Ono et al., 1986; Li et al., 2013). In addition to caloric value, sucrose’s palatability is the affective or hedonic attribute of sweetness that determines whether to like it or not (Grill and Berridge KC, 1985). Despite the importance of palatability to promote overconsumption, the specific LHA cell-type(s) identity involved in processing sucrose’s palatability has remained elusive. Our results demonstrated that a subpopulation of LHA^Vgat+^ GABAergic neurons encodes sucrose’s palatability by exhibiting two opposite modulatory patterns, either correlating positively or negatively with the palatability index, with a bias towards a positive correlation. Furthermore, opto-stimulation of LHA^Vgat+^ cell somas promoted the approach and intake of the most salient available and palatable tastant. In contrast, opto-self-stimulation promoted increased liquid intake of the less-attractive and proximal stimuli, despite having more palatable but distal tastants available. These findings show that LHA^Vgat+^ neurons compute and/or combine, at least, two types of information: one related to stimulus proximity and the other to palatability that results in enhancing stimulus-saliency (Nieh et al., 2016). Experiments with solid food also unveiled that transgenic mice spent more time near the most palatable food available. More important, among the many other functions already ascribed to these neurons (see below and Nieh et al., 2016), our data uncovered a new function of LHA^Vgat+^ neurons as physiological potentiators of sucrose-induced oromotor palatability responses.

Previous studies have shown that LHA^Vgat+^ population contains many subpopulations with different functional responses, with at least one ensemble responding to appetitive (approach) and others to consummatory behaviors (Jennings et al., 2015). Furthermore, activation of these neurons is rewarding, induces voracious eating (Jennings et al., 2015; Navarro et al., 2016), and promotes interaction of the nearest stimulus (either objects or other mice) (Nieh et al., 2016), suggesting that they play a role in multiple motivated behaviors. However, it is also known that LHA connects and receives direct inputs from multiple cortical and subcortical gustatory regions (Simerly, 2004; Berthoud and Münzberg, 2011), and some electrophysiological studies report that LHA neurons respond to gustatory stimuli – in particular to tastant palatability (Norgren, 1970; Ono et al., 1986; Schwartzbaum, 1988; Yamamoto et al., 1989; Karádi et al., 1992). As noted in rodents, palatability is operationally defined as the enhancement of hedonically positive oromotor responses induced by stimulating the tongue with ascending sucrose concentrations (Berridge and Grill, 1983; Spector et al., 1998; Villavicencio et al., 2018). Specifically, these hedonically positive oromotor responses may include an increase in the lick rate or the bout size. In agreement with this definition, we found that a subpopulation of LHA palatability related-neurons tracked licking oromotor-related responses by increasing or decreasing their activity in a sucrose-concentration dependent manner. Within a session, LHA neurons tracked the sucrose’s palatability rather than satiety or hunger signals. We showed that these LHA^Vgat+^ neurons could function as enhancers of sucrose’s palatability. Optogenetic activation of these neurons can also enhance water’s palatability if it is the nearest stimulus. We found an increased lick rate for water during these neurons’ activation as if the animal were sampling a high sucrose concentration. Moreover, their activation promotes the intake of liquid sucrose (or solid granulated sugar cube). These neurons also increased the consumption of other more palatable stimuli like high-fat pellets (see Supplementary Figure 5; Video 2), similar to the other GABAergic neurons but in zona incerta (Zhang and van den Pol, 2017). Thus, our data demonstrate that activation of LHA GABAergic drives the intake of the most palatable stimulus available in the animal’s environment.

Although activation of LHA^Vgat+^ promotes substantial feeding behavior, it has become clear that LHA^Vgat+^ neurons are not directly involved in evoking hunger (Burnett et al., 2016; Navarro et al., 2016; Marino et al., 2020), as AgRP neurons in the arcuate do (Chen et al., 2016). In this regard, and unlike AgRP neurons, pre-stimulation of LHA GABAergic neurons did not trigger a sustained sucrose intake in the absence of continuous activation (Supplementary Figure 4). Thus, to induce a consummatory behavior, these neurons are required to remain active. Moreover, and in agreement with these findings, we found that the intake induced by LHA^Vgat+^ neurons convey a positive valence signal that combines both stimulus proximity and palatability related information. Thus, these neurons enhance the saliency of nearby hedonically positive stimuli, whether those stimuli are sapid chemicals, as we show, or social cues, as in the approach behavior towards juvenile or female intruders and new objects (Nieh et al., 2016).

It is important to highlight that opto-self-activation of LHA^Vgat+^ neurons resembles many hallmark behaviors evoked by LHA electrical stimulation. In particular, our results could shed some light on why, at low-intensity electrical currents, lever pressing to deliver ICSs only occurs if food (or sucrose) is in close proximity (Mendelson, 1967; Coons and Cruce, 1968; Valenstein et al., 1968; Valenstein and Phillips, 1970). Consistent with its role in enhancing sucrose’s palatability, a subpopulation of GABAergic neurons could indirectly explain why sweet tastants further potentiate the rate of LHA electrical ICSs (Poschel, 1968). We concluded that a subpopulation of LHA^Vgat+^ neurons could account for many, if not all, of these electrically induced phenomena. Also, we found differences between unspecific LHA stimulation and our targeted LHA^Vgat+^ stimulation. Unlike electrical LHA stimulation, we found that the laser-bound feeding was observed in all tested VGAT-ChR2 mice (n = 31, Figures 4–12). In contrast, to the high variability found in rats exhibiting LHA electrically-induced feeding, one study reported that only 12 of 34 rats showed stimulus-bound feeding (Valenstein and Cox, 1970). Similarly, a large variability was observed when unspecific bulk optogenetic stimulation activated *all* cell-types found in LHA, simultaneously (Urstadt and Berridge, 2020). These studies reported that after repeated LHA stimulation (either electrically or with optogenetics), some subjects switched from exhibiting stimulus-bound feeding to only self-stimulating, suggesting that these two processes were flexible and not correlated (Gigante et al., 2016; Urstadt and Berridge, 2020). In contrast, we found that repeated stimulation of LHA^Vgat+^ neurons increases laser-induced feeding (licking). Likewise, the correlation between optogenetic self-stimulation and laser-bound feeding increases through stimulation days (Figure 8). That is, the more the animals self-stimulated, the stronger the evoked laser-bound licking was. Thus, LHA^Vgat+^ neurons are the common neural substrate for evoking both feeding and reward. Though, it was recently shown that they do it by using two projection pathways; reward via a VTA projection and feeding via the peri-Locus Coeruleus nuclei (Marino et al., 2020).

Given that LHA is involved in reward and aversion (Ono et al., 1986), we next tested whether LHA^Vgat+^ neurons could promote bitter tastants’ intake. We found that opto-stimulation of LHA^Vgat+^ neurons failed to promote quinine intake, a bitter tastant, or tolerance of an aversive airpuff when sucrose was also available. Thus, these neurons play a minimal role in increasing the preference for a proximal but aversive stimulus over distal sucrose. Furthermore, in sated mice and using a single bottle test, these neurons also failed to increase quinine intake. Unexpectedly, during water deprivation, a copious quinine consumption was observed. These results demonstrate that the consummatory drive induced by the activation of GABAergic neurons largely depends on the palatability of stimulus and animal’s internal state. These results agree with previous findings that chemogenetic inhibition of LHA^Vgat+^ neurons did not alter the quinine rejection responses. Thus, in sated mice, these neurons are not necessary to express hedonically negative responses induced by bitter tastants (Fu et al., 2019). However, they did not explore their sufficiency. Our results showed that water deprivation temporarily gated LHA^Vgat+^ neurons to promote quinine intake and further demonstrated that their activation is sufficient to increase acceptance of an aversive tastant during water deprivation.

The LHA comprises multiple heterogeneous and overlapping populations based on the expression of genetic markers for neuropeptides, receptors, and proteins involved in the synthesis and vesicular packaging of neurotransmitters (Bonnavion et al., 2016; Mickelsen et al., 2019). Thus, the LHA^Vgat+^ population can be further subdivided into GABA neurons expressing leptin receptor (LepRb) (Leinninger et al., 2009; Mickelsen et al., 2019), or the neuropeptide galanin (Gal), or neurotensin (Nts) (Qualls-Creekmore et al., 2017; Kurt et al., 2019; Mickelsen et al., 2019). It is known that the activation of LHA-GABA-LepRb is rewarding (Giardino et al., 2018), similar to LHA^Vgat+^ neurons. Likewise, LHA GABA-Gal expressing neurons are related to food reward behavior, but unlike LHA^Vgat+^, these neurons do not promote food consumption (Qualls-Creekmore et al., 2017). From these subpopulations, only the LHA GABA-Nts neurons recapitulate some (but not all) of the behavioral effects reported here. Unlike LHA^Vgat+^ neurons, a previous study found that LHA GABA-Nts neurons do not increase chow intake. Instead, they promote the liquid intake of palatable tastants (water, NaCl, and sucrose). In sated mice, chemogenetic activation of GABA-Nts increased the intake of bitter quinine, albeit with lower magnitude, when it was the only liquid available to eat with chow food, a marked contrast to our findings with broad LHA GABAergic neurons activation. Although they did not explore quinine intake in the absence of chow food or water-deprived mice. However, they performed a two-bottle test and found that GABA-Nts neurons increased water intake over quinine, suggesting that activation of these neurons is not involved in driving mice’s preference for bitter tastants, similar to what we found for the LHA^Vgat+^ population. Also, similar to our findings, activation of LHA GABA-Nts induced water-drinking, which was further facilitated if the solution was sucrose (Kurt et al., 2019). Thus, it will be interesting to determine the role that LHA GABA-Nts neurons play in encoding and potentiating sucrose’s palatability. It follows that the LHA contains nested functions encoded in each subpopulation (or cell-types) that are then recruited selectively to exert a more refined control over feeding and reward.

A caveat of this study is that we employed bacterial artificial chromosomes (BAC) transgenic strain mouse, VGAT-ChR2, that constitutively expressed ChR2 in GABAergic neurons expressing the gene for the VGAT (Zhao et al., 2011). In this model, we cannot rule out the unintended activation of GABAergic terminals from distal regions (Thoeni et al., 2020), which also occurs with classic electrical stimulation. Nevertheless, in the more specific transgenic model, the Vgat-ires-Cre mice have found a similar feeding-bound behavior for chow food (Marino et al., 2020), as we have shown here (see Chow in Figures 9B and C; and Video 5). Moreover, the VGAT-ChR2 transgenic model affords important advantages such as a consistent expression of ChR2 (Zeng and Madisen, 2012) and heritable transgene expression patterns across experimental cohorts (Ting and Feng, 2013), which increased reproducibility across animals tested. It is also a more selective model to characterize GABAergic neurons (excluding the glutamatergic component) and their effects recapitulating classical effects observed with electrical LHA stimulation (Delgado and Anand, 1953; Phillips and Mogenson, 1968).

In summary, here we found that at least a subpopulation of LHA^Vgat+^ neurons could be an important hub that links stimulus-proximity and palatability related information to potentiate the palatability of nearby energy-rich foods, especially those containing sucrose.

## Author Contributions

A.G., A.C., J. LI., and R.G. designed research; A.G., A.C., J. LI., L.PS., and M.L. performed research; M.V. contributed unpublished analytic tools; A.G., A.C., J. LI., and R.G. analyzed data; A.G. and R.G. wrote the paper; all authors reviewed and approved the manuscript.

## Acknowledgments

Aketzali Garcia had a CONACyT doctoral fellowship, and data in this work is part of her doctoral dissertation in the Posgrado en Ciencias Biomédicas of the Universidad Nacional Autónoma de México. We want to thank Martin Vignovich and Professor Sidney A. Simon for helpful comments in an early version of this manuscript. We thank Mario Gil Moreno for building homemade optrode and valuable help of Juan de Dios Rodriguez-Callejas during the acquisition of confocal microscope images. We also thank Ricardo Gaxiola, Victor Manuel García Gómez, and Fabiola Hernandez Olvera for invaluable animal care.

This manuscript has been released as a pre-print at Biorxiv, (Garcia et al., 2020).

