## Supplementary figures for "Lateral Hypothalamic GABAergic neurons encode and potentiate sucrose’s palatability"

### *Supplementary Material*

#### Supplementary Figures

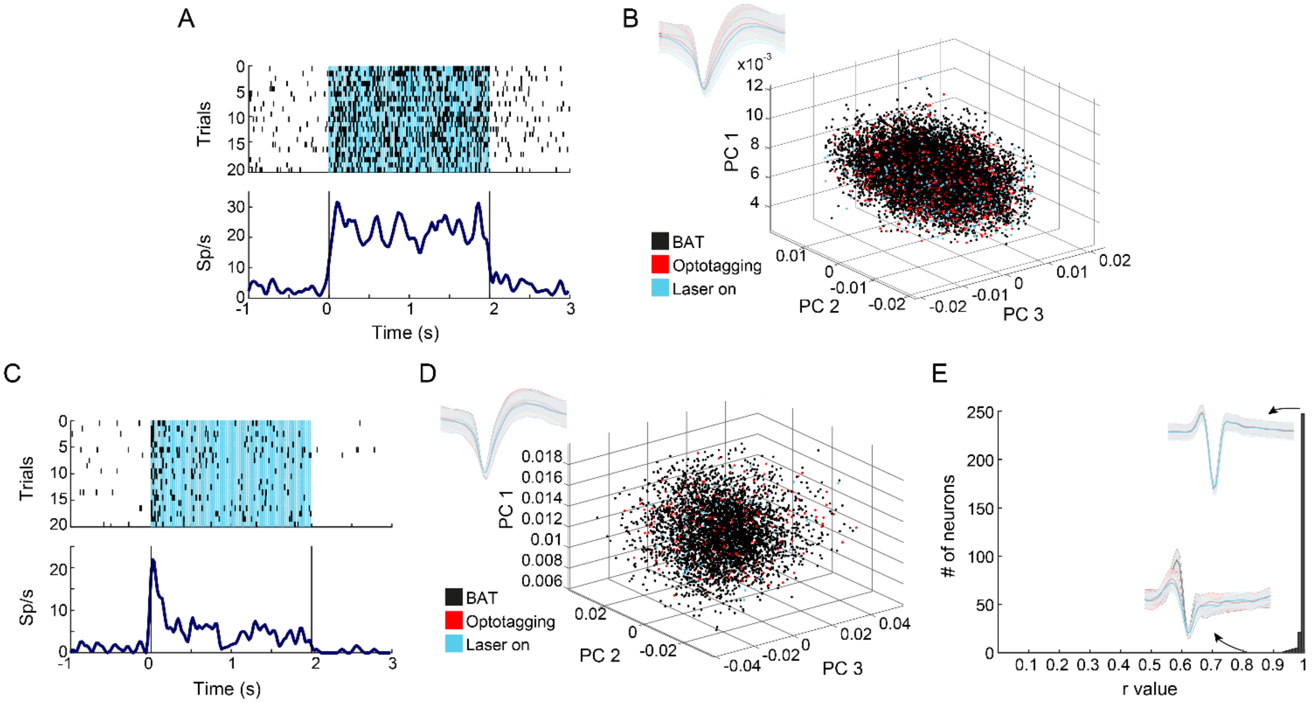


**Supplementary Figure 1.** Waveform stability in extracellular recordings of LHA neurons across the brief-access test (BAT) and the optotagging session. **(A)** Representative raster plot of a LHA$^{Vgat+}$ neuron recorded during the optotagging session. Spiking responses were aligned (Time = 0 s) to the first laser pulse. Black ticks depict action potentials and blue marks laser pulses (at 50 Hz). Below, PSTHs (Spikes /s, Sp/s). Vertical black lines indicate laser onset (Time = 0 s) and offset (Time = 2 s), respectively. **(B)** *Inset upper left* depicts average waveforms of a single neuron showed in A across the BAT (black), the optotagging stimulation (red), and when the laser was turned “on” (blue). Right, each dot represents an action potential during the entire session (same color-coded). Values are plotted in a three-dimensional space computed with the first three principal components (PCs) of the waveforms. **(C-D)** Same conventions as in (A-B). ***E*****,** Histogram of the Pearson correlation coefficients of waveforms recorded during BAT against those recorded in the optotagging session. Waveforms depict two examples, one depicting the worst and the other the best correlation included in analysis.

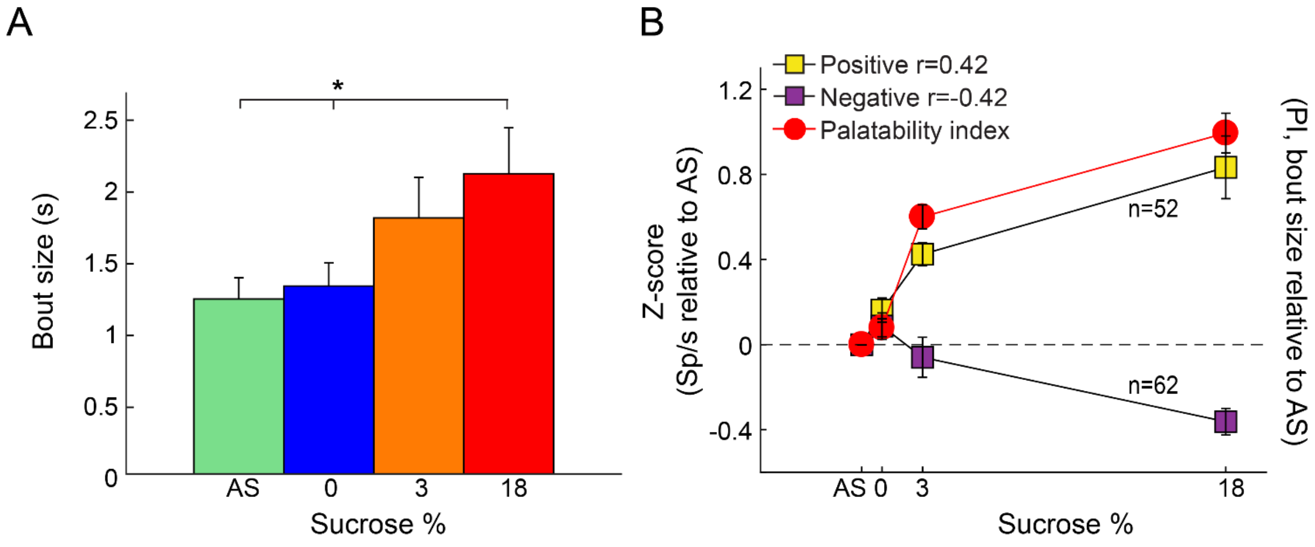


**Supplementary Figure 2.** Palatability-related neurons identified by using the bout size to compute the Palatability Index **(A)** Average bout size during the entire reward epoch. Bout size of sucrose at 18% is statistically higher (p <0.05) than AS and water (0). Using one-way ANOVA followed by the Holm Sidak test. **(B)** Z-score normalized activity (relative to AS trials) for LHA neurons with either positive (yellow) or negative (purple) correlation against PI (red circles).

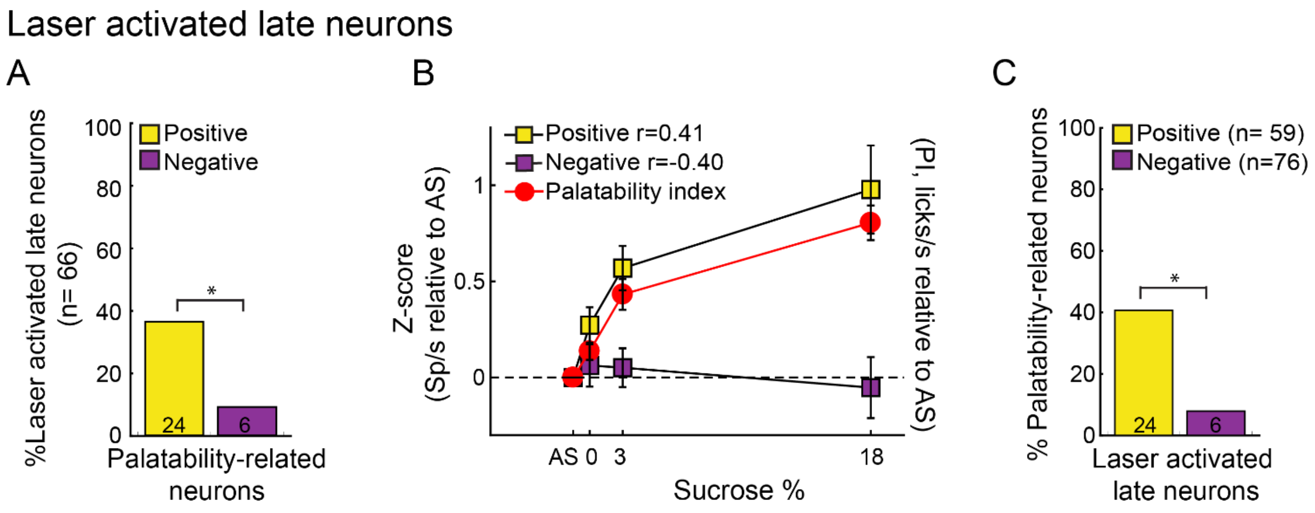


**Supplementary Figure 3.** Laser-activated late neurons encode sucrose's palatability **(A)** Late neurons were activated > 15 ms after the laser onset. Percentage of late neurons whose activity correlates positively or negatively correlates with sucrose's PI. * $p$ < 0.001, chi-square. **(C)** Z-score activity (relative to AS) for late neurons and its correlation with the sucrose's PI. **(D)** Percentage of Positive or Negative laser-activated late palatability-related neurons. * $p$ < 0.0001, chi-square.

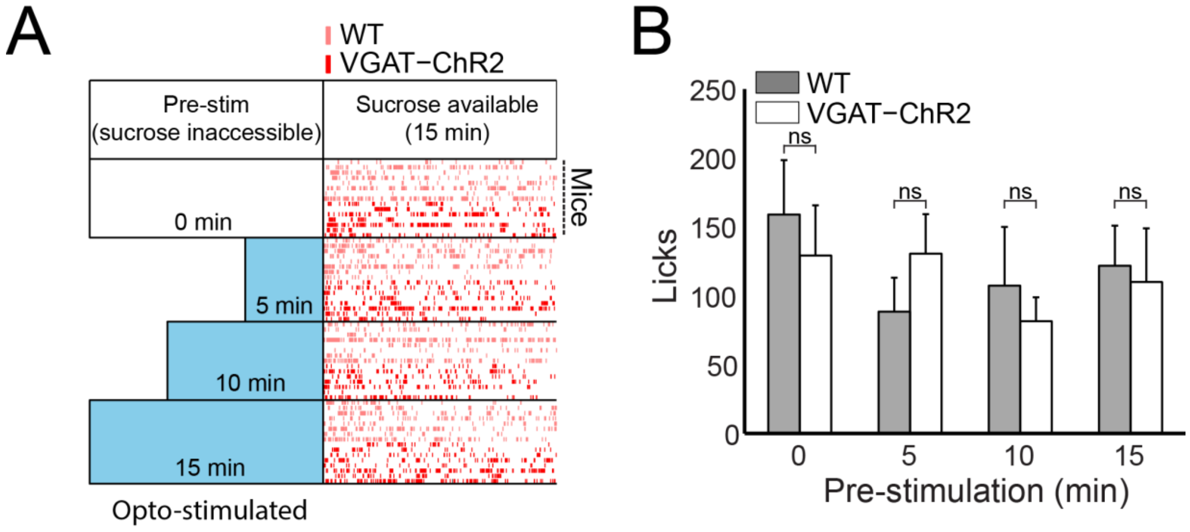


**Supplementary Figure 4.** Pre-stimulation of LHA$^{VGAT+}$ neurons did not trigger a "hunger state" in that a sustained consummatory behavior is not observed. **(A)** Schematic of the pre-stimulation protocol and raster plot from VGAT-ChR2 (licks were plotted as red ticks) and WT (licks as light red ticks) mice. For this experiment, all mice underwent all pre-stimulation protocols following a Latin square design: 1) mice were pre-stimulated for either 0, 5, 10, or 15 min with no sucrose available. After pre-stimulation, mice received 15 min Free access to a sipper filled with sucrose 10%. **(B)** The average number of licks for sucrose as a function of pre-stimulation.

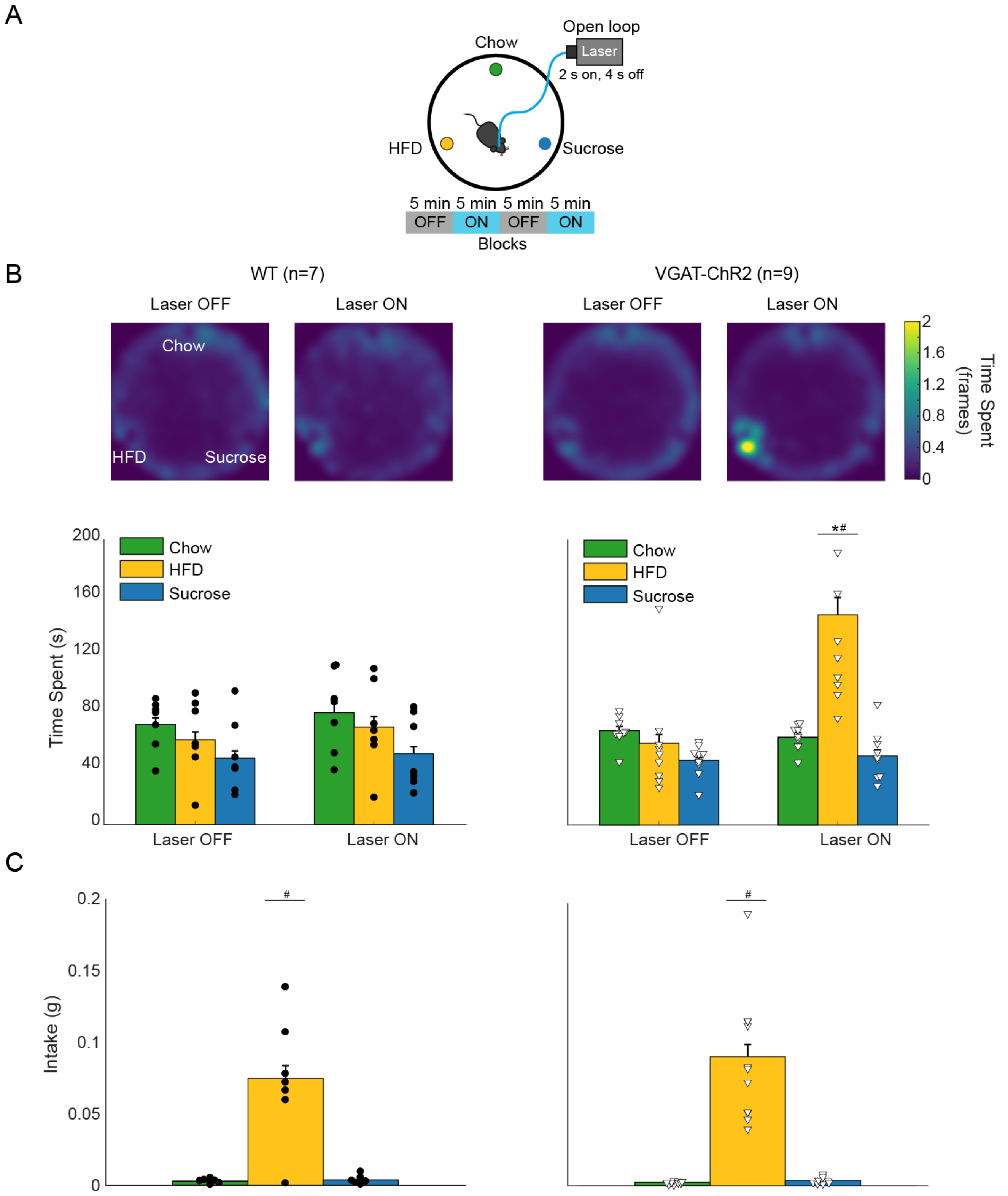


**Supplementary Figure 5.** Activation of LHA$^{Vgat+}$ neurons increases the time spent near the most palatable High Fat Diet (HFD). **(A)** In a circular arena, three plates containing Chow (green), HFD (yellow), and granulated sucrose cube (blue) were available simultaneously. In a 20 min session, the laser was turned "on" following an open loop protocol (2 s on 4 s off during 5 min block with laser and 5 min with no-laser; see Figure 5A). **(B)** Average heat map and time spent in the open field. To create each heat map, we identified the centroid of the mouse at every video frame. Colorbar indicates the number of frames the subject was detected in each pixel; higher values indicate the mice stayed in that place for a longer time. Time spent (below) was calculated as the distance between the mouse's centroid relative to each food plate. We quantify a food interaction when the subject was within 60 mm distance relative to the plate's center and remained there for at least 1 s. Each dot and triangle represent a single individual. **(C)** Intake of Chow, HFD, and granulated sucrose cube. * Denote statistically significant differences ($p < 0.0001$) relative to WT. # $p < 0.0001$ between a stimulus relative to the others. One-way ANOVA followed by the Holm Sidak test. See Video 2.

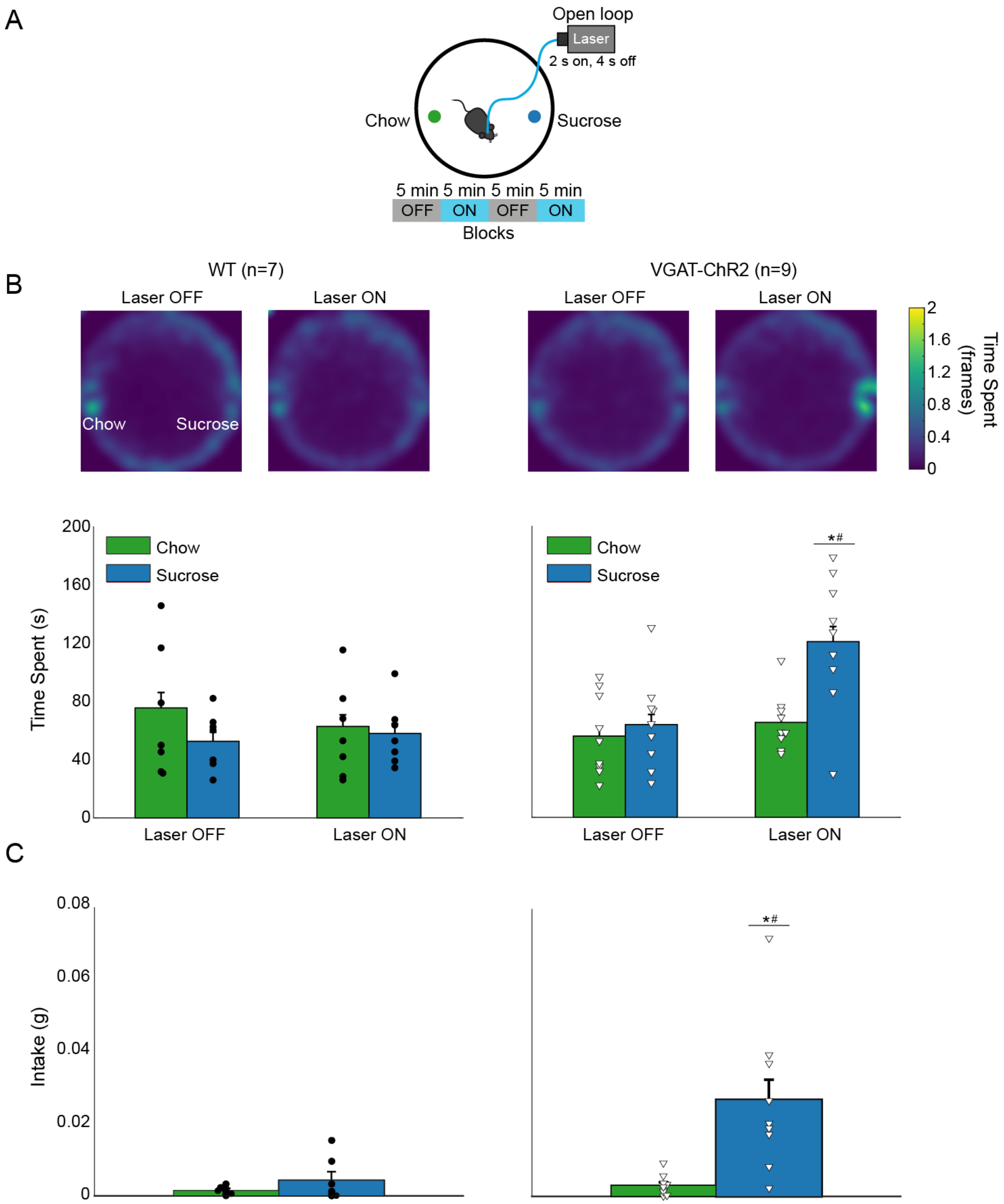


**Supplementary Figure 6.** Activation of LHA$^{Vgat+}$ neurons promotes intake to the most palatable solid food. ***(A)*** In a circular arena, Chow pellet and granulated sucrose cube were presented simultaneously.

The laser was turned "on" in an open loop protocol. **(B)** Mean heat map and time spent. The same conventions are in Supplementary Figure 5. **(C)** Intake of Chow and granulated sucrose cube. * Statistically significant differences ($p < 0.0001$) relative to WT. # $p < 0.0001$ difference between a stimulus relative to the others. One-way ANOVA followed by the Holm Sidak test. See Video 3.

**Video 1. Open loop activation of LHA$^{Vgat+}$ neurons.** Water, 3 wt%, and 18 wt% sucrose was delivered in each port. Opto-stimulation occurred in 5 min block of no-laser and 5 min block with the laser.

**Video 2. Open loop activation of LHA$^{Vgat+}$ neurons promotes preference toward HFD over granulated sucrose and Chow pellet**. At equidistant position, three foods were placed on a small plate, and each contained either HFD, Chow food, or granulated sucrose cube. Opto-stimulation occurred in 5 min block of no-laser and 5 min block of the laser for 20 min.

**Video 3. VGAT-ChR2 mice prefer granulated sucrose over Chow food.** Preference and intake of granulated sucrose cube vs. Chow pellet after open-loop activation of LHA GABAergic neurons.

**Video 4. Closed-loop activation of LHA$^{Vgat+}$ neurons.** Mice opto-self-stimulate in each head entry in the central licking port (where water is the nearest stimulus). Some VGAT-ChR2 mice exhibited a preference for the most palatable stimulus available (sucrose 18%), whereas others preferred the nearest but less palatable stimulus (water) over sucrose 18%

**Video 5. Real-time place preference in an open field task.** At equidistant separation, four stimuli were placed on a small plate, and each contained either wood cork, Chow pellet, chocolate pellets, or a sipper a 10% sucrose solution. Mice were photostimulated by 2 s on and 4s off when crossing the designated area.

**Video 6. In some VGAT-ChR2 mice, activation of LHA$^{Vgat+}$ neurons induces gnawing to a cork**. Wood cork was placed in a transparent box, whereas mice were opto-stimulated in an open loop protocol.
